# Downregulation of LATS1/2 Drives Endothelial Senescence-Associated Stemness (SAS) and Atherothrombotic Lesion Formation

**DOI:** 10.1101/2025.06.19.660635

**Authors:** Sivareddy Kotla, Jonghae Lee, Kyung Ae Ko, Weiqing Chen, Venkata Subrahmanya Kumar Samanthapudi, Oanh Hoang, Gilbert F. Mejia, Shengyu Li, Keri L. Schadler, Luis Antonio Rivera, Masaki Imanishi, Kay Carlene Tavares Samperio, Jung Hyun Kim, Kelia C. Ostos-Mendoza, Karla N. Mariscal-Reyes, Anita Deswal, John P. Cooke, Keigi Fujiwara, Nicolas L. Palaskas, Efstratios Koutroumpakis, Young Jin Gi, Rajneesh Pathania, Criag Morrell, Philip L. Lorenzi, Lin Tan, Iqbal Madhmud, Nordin M.J. Hanssen, Laurent Yvan-Charvet, Eduardo N. Chini, Joerg Herrmann, Hernan G. Vasquez, Ying H. Shen, James F. Martin, Haodong Xu, Erin H Seeley, Jared K. Burks, Paul S. Brookes, Guangyu Wang, Nhat-Tu Le, Jun-ichi Abe

## Abstract

**Background:** Atherothrombosis, the main event leading to acute coronary syndrome (ACS), is strongly linked to disturbed blood flow (d-flow) regions. Although the involvement of the Hippo pathway and its kinases Large Tumor Suppressor Kinase 1and 2 (LATS 1 and 2) in mechanical stress responses is known, the mechanisms by which d-flow simultaneously induces senescence, proliferation, and atherothrombosis remain unclear.

**Methods:** The role of endothelial cells (EC)-specific LATS1/2 was examined using EC specific knock-out (EKO) mice in a partial left carotid ligation (PLCL) model. Plaque spatial multi-omics analysis was performed by integrating imaging mass cytometry, sequential immunofluorescence (COMET™), and spatial metabolomics at the single-cell level in human and mouse atherosclerotic plaques.

**Results:** In tamoxifen-inducible *Lats1^homo(^*^-/-)^ /*Lats2 ^homo(^*^-/-)^ EC-specific knockout (EKO) mice, deletion of LATS1/2 induced by tamoxifen led to fatal outcomes, characterized by severe systemic edema and markedly increased vascular permeability. In contrast, *Lats1^het(+^*^/-)^/*Lats2 homo(*-/-)-EKO mice survived and developed atherothrombotic plaques exhibiting neovascularization even without further additional dietary or genetic intervention. Spatial proteomics analysis revealed that LATS1/2 depletion in ECs triggered a senescence-associated stemness (SAS) phenotype, primarily driven by CD38 upregulation. Complementary spatial metabolomics profiling demonstrated a significant increase in sulfite and taurine within LATS1/2-deficient plaques, indicating lowered sulfite oxidase (SUOX) activity. Mechanistically, CD38 upregulation was found to suppress SUOX expression, induce the reverse mode of mitochondrial complex V, and increase succinate dehydrogenase (SDH) activity along with ATP consumption. Paradoxically, despite ATP depletion, this metabolic disturbance enhanced glutamate metabolism and the tricarboxylic acid (TCA) cycle, sustaining EC proliferation under energetically stressed conditions. The combined effect of LATS1/2 deletion and CD38 activation established a unique EC phenotype defined by increased SAS, leading to proliferation, senescence, and eventual cell death. These pathological processes culminated in the formation of atherothrombotic plaques, which were attenuated by inhibition of CD38. Notably, a similar phenotype—marked by metabolically active ECs—was observed in human atherothrombotic plaques, suggesting translational relevance.

**Conclusion:** Loss of LATS1/2 in ECs induces SAS state that promotes excessive EC proliferation, senescent cell accumulation, and the development of structurally fragile, leaky neo vessels—hallmarks of atherothrombotic lesions. CD38-mediated SUOX deficiency further amplifies this pathological process by inducing mitochondrial dysfunction, depleting ATP, and triggering compensatory upregulation of glutamate and TCA cycle metabolism. These findings identify a novel LATS1/2–CD38–SUOX axis in ECs that orchestrates SAS-driven atherothrombosis. Targeting CD38 may represent a promising therapeutic strategy to mitigate vascular dysfunction and plaque instability in high-risk ACS patients.

**Graphical abstract:** 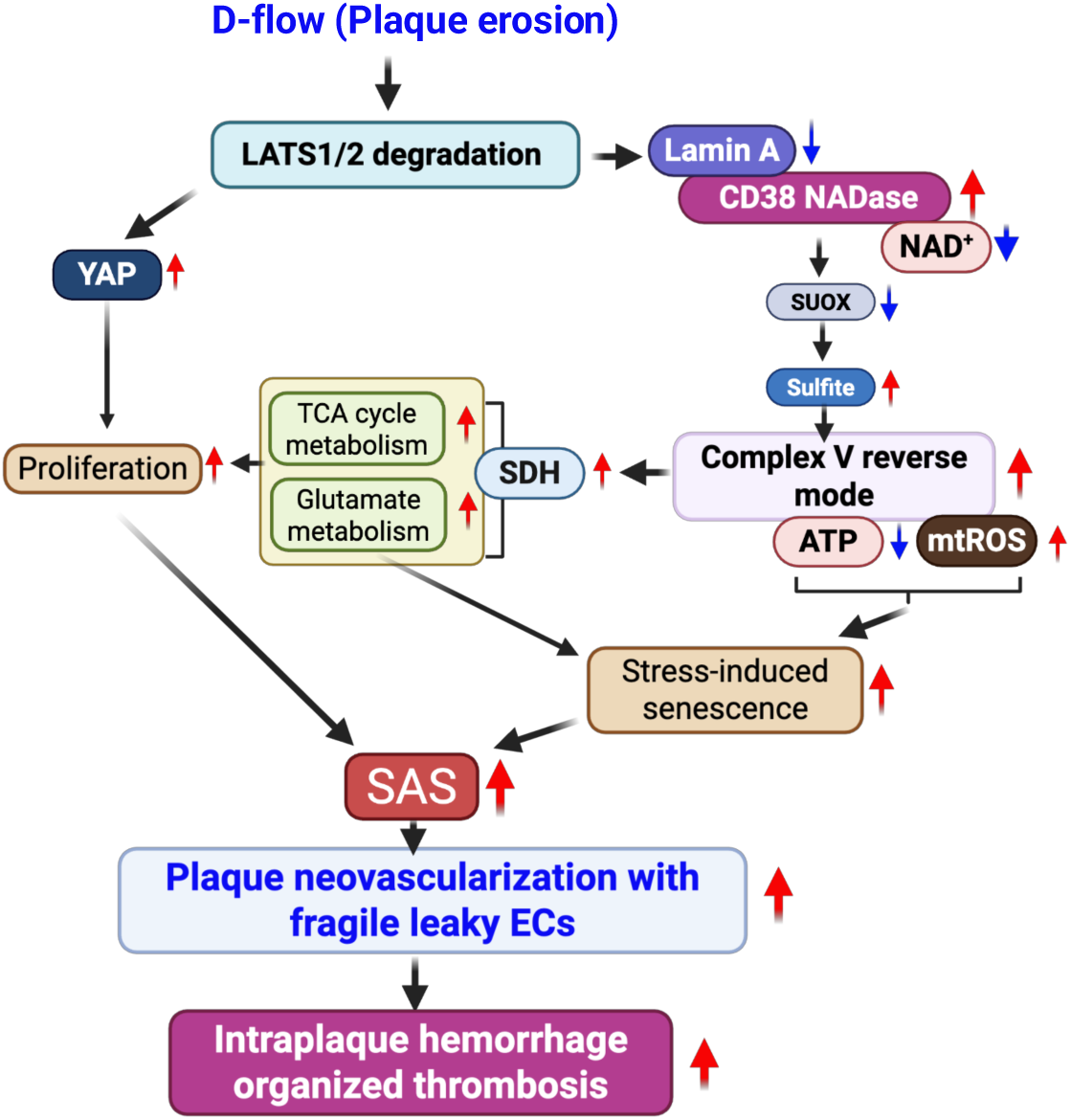

Under normal physiological conditions, LATS1/2 and Lamin A work together to suppress CD38 expression. Lamin A binds directly to the CD38 promoter to repress transcription, and LATS1/2 interact with Lamin A to reinforce this suppression. This collaboration helps maintain low CD38 activity and preserves cellular NAD⁺ levels. However, under disturbed flow (d-flow), both LATS1/2 and Lamin A are downregulated. The loss of this dual repression leads to increased CD38 NADase expression. Elevated CD38 accelerates NAD⁺ consumption, causing NAD⁺ depletion—a hallmark of cellular senescence. Reduced NAD⁺ disrupts key metabolic and stress-response pathways, contributing to the onset of the senescent state. At the same time, CD38 suppresses sulfite oxidase (SUOX), leading to sulfite accumulation and mitochondrial redox imbalance. This shift activates the reverse mode of mitochondrial Complex V, which decreases ATP production and increases mitochondrial ROS, intensifying metabolic and oxidative stress in endothelial cells (ECs). In response, ECs compensate by upregulating succinate dehydrogenase (SDH), enhancing TCA cycle activity and glutamate metabolism. This metabolic adaptation provides the biosynthetic building blocks needed for cell growth and proliferation. As a result, ECs adopt a paradoxical phenotype: they show classical features of stress-induced senescence (such as NAD⁺ depletion, oxidative stress, and cell cycle arrest signals), while simultaneously undergoing metabolic activation and proliferation, also mediated by YAP. This defines a non-canonical endothelial program known as senescence-associated stemness (SAS), characterized by the formation of abnormal, proliferative, yet fragile neovessels. These dysfunctional vessels contribute to atherothrombosis, setting this process apart from the more stable lesions typical of conventional atherosclerosis.

## INTRODUCTION

Plaque rupture and plaque erosion account for the great majority of acute coronary syndromes; the remainder being attributed to intraplaque hemorrhage and calcified nodules^1,2^. Disturbed (d-) flow conditions have been indicated to play a mechanistic role in the destabilization of atherosclerotic plaques^3,4^. The contribution of both EC proliferation and senescence in plaque destabilization and atherothrombotic lesions has been suggested^5,6^, but how both EC proliferation and senescence can be induced simultaneously by d-flow has not been well studied because of a lack of proper animal models of plaque rupture and erosion.

The Hippo pathway emerges as a key regulator in this context due to its central role in mechanotransduction and endothelial responses to hemodynamic forces^7,8^. At its core, the mammalian Hippo pathway consists of two kinase pairs: MST1/2 and LATS1/2. MST1/2, together with the adaptor protein WW45, phosphorylate and activate LATS1/2, a process further facilitated by MOB1, which enhances LATS1/2 activation through direct interaction^9,10^. Activated LATS1/2 in turn phosphorylate YAP at serine 127, promoting its cytoplasmic retention via 14-3-3 binding and suppressing its transcriptional activity^9,10^. When dephosphorylated, YAP/TAZ translocate into the nucleus, bind TEAD, and drive gene expression programs that promote cell proliferation^11^, inhibit apoptosis, and regulate organ size^9,12,13^.

Importantly, YAP/TAZ activity is strongly influenced by mechanical cues beyond canonical MST1/2 signaling, including changes in cell morphology and cytoskeletal tension. In ECs, d-flow promotes YAP/TAZ nuclear translocation, enhancing pro-inflammatory gene expression, while laminar flow (l-flow) suppresses YAP activation through mechanisms such as increased phosphorylation^8^ or SIRT1-mediated deacetylation^14^. These findings underscore the sensitivity of YAP/TAZ to hemodynamic forces. However, the upstream regulation of YAP/TAZ by LATS1/2 in response to flow, particularly under d-flow conditions, remains poorly understood, highlighting a critical gap in our understanding and positioning LATS1/2 as important mediators of flow-sensitive endothelial signaling.

In addition to its role in mechanotransduction, YAP/TAZ also regulates cellular senescence in a context- and cell-type-dependent manner. In stromal cells, YAP/TAZ prevent senescence by preserving nuclear envelope integrity and suppressing the cGAS-STING pathway^15^. Conversely, in aging vascular tissues, YAP is often upregulated, and its inhibition has been shown to reduce senescence—likely through restoration of autophagic flux and attenuation of mTOR signaling^16^. LATS2 contributes to senescence independently by repressing the DREAM complex and E2F target genes, potentially through a feedback loop with YAP^17^. Moreover, LATS2 overexpression alone can induce cell cycle arrest and upregulate senescence markers in epithelial cells^17^. Although these findings suggest that LATS1/2 may participate in regulating endothelial senescence, their role in d-flow-induced endothelial senescence remains to be elucidated^18^.

In this study, we assess the role of endothelial LATS1/2 in inducing senescence, proliferation, inflammation, and metabolic remodeling in vivo and in vitro. We use EC-specific LATS1 and LATS2 knockout mice (EKO), employing a novel combination of COMET™ (multi-fluorescence imaging) and imaging mass cytometry (IMC) techniques^19^, along with spatial metabolomic analysis and single-cell evaluation by integrating spatial proteomics and metabolomics data. Our findings reveal that the depletion of LATS1/2 induces SAS, supported by a unique metabolomic shift in ECs, leading to atherothrombosis with neoangiogenesis formation.

## METHODS

Details of the antibodies and reagents, experimental models and subject details, experimental procedures, metabolomics analysis including carbon-13 tracing experiments for metabolic flux analysis, imaging with COMET™ multi-fluorescence imaging and imaging mass cytometry (IMC), spatial metabolomics analysis, and power calculation are included in the online supplemental information.

### Data Sharing Availability

The data, analytic methods, and study materials that support the findings of this study are available in the Data supplement or from the corresponding authors upon reasonable request.

### Institutional Approvals

Human aortic tissues were obtained as de-identified biospecimens in compliance with protocols approved by the Institutional Review Board (IRB) at Baylor College of Medicine (IRB protocol H-12515), which governs the ethical collection and de-identification of human cardiovascular specimens. The University of Texas MD Anderson Cancer Center reviewed the use of these de-identified tissues and, under IRB protocol 2023-1363, determined that their use does not constitute human subjects research. Consequently, no additional IRB approval or informed consent was required for experimental use. All procedures adhered to applicable federal regulations, including the Health Insurance Portability and Accountability Act (HIPAA) and the Common Rule (45 CFR 46).

All animal studies were conducted in accordance with institutional and federal ethical guidelines and were approved by the Institutional Animal Care and Use Committees (IACUC) at both the Texas A&M Institute of Biosciences and Technology (protocols 2014-0231 and 2017-0154) and The University of Texas MD Anderson Cancer Center (protocols 00001652 and 00001952).

### Statistical Analysis

All statistical analyses were performed using GraphPad Prism software version 9.0.0 or 10 (GraphPad Software, San Diego, CA, USA). The Shapiro–Wilk test was used to assess the normality of each dataset. For comparisons among more than two groups, ordinary one-way or two-way analysis of variance (ANOVA) was conducted, followed by Tukey’s post hoc test, provided the data met normality assumptions. For two-group comparisons with normally distributed data, unpaired two-tailed Student’s *t*-tests were applied. When data failed the Shapiro–Wilk normality test, the nonparametric Mann–Whitney *U* test was used for two-group comparisons, and the Kruskal–Wallis test followed by Dunn’s post hoc test was applied for multi-group comparisons. A *P* value < 0.05 was considered statistically significant, except for analyses involving Imaging Mass Cytometry (IMC) data. All experiments were performed independently unless otherwise indicated.

All data visualizations, including violin plots and bar graphs, were generated using GraphPad Prism version 10 (GraphPad Software, San Diego, CA, USA). Detailed statistical information— including sample sizes (*n*), normality test outcomes, statistical methods used, and *P* values for each figure—is provided in Table S1.

### IMC and COMET Data Analysis

Image files from two distinct platforms—COMET (a sequential immunofluorescence platform) and Imaging Mass Cytometry (IMC) using the Fluidigm Hyperion system—were exported as OME-TIFF and MCD (mass cytometry data) formats, respectively. These files were imported into Visiopharm software and aligned using the TissueAlign module with a U-net-based approach to generate a unified image database for high-dimensional single-cell analysis.

Co-registration of COMET and IMC images was performed using shared marker channels (CD4, CD8, CD31, and αSMA) to enable precise spatial alignment across modalities. Cell segmentation was carried out using a protocol trained on DAPI+ nuclear signals from COMET images, followed by a dilation step to define accurate cell boundaries. A phenotyping pipeline was then applied within the COMET dataset to assess the expression of functional and lineage-specific markers. Marker intensity thresholds were used to classify distinct cell populations, including those exhibiting endothelial cell (EC)-like characteristics.

All data visualizations, including violin plots and bar graphs, were generated using GraphPad Prism version 10 (GraphPad Software, San Diego, CA, USA). Detailed statistical information— including sample sizes (*n*), normality test outcomes, statistical methods used, and *P* values for each figure—is provided in Table S1.

**Supplementary Table 1:** Statistical analysis for each experiments

**Supplementary Table 2:** Controls for each relative value:

**Supplementary Table 3:** Antibodies for IMC and COMET

**Supplementary File 1:** Spatial proteomics intensity data used in Fig.S2 and Fig.3A in mouse plaque study. Immune cells and ECs, and two ROIs (surface and plaque) data were included.

**Supplementary File 2:** Spatial metabolomics data used in Figure 5 in mouse plaque study. CD31+ ECs and two ROIs (surface and plaque) data were included.

**Supplementary File 3:** Spatial proteomics data used in human plaque study used in Fig.7.

**Supplementary File 4:** Spatial metabolomics data used in human plaque study used in Fig. 8.

**Supplementary File 5:** Glutamine tracing data used in Fig.6.

**Supplementary File 6:** Glucose tracing data used in Fig.6.

## Results

### Endothelial LATS1/2 are indispensable for vascular barrier maintenance and survival

To elucidate the physiological role of LATS1/2 in vascular homeostasis, we generated tamoxifen-inducible endothelial-specific homozygous knockout mice (*Lats1^homo(-/-)^/Lats2^homo(-/-)^*), hereafter *Lats1*^-/-^/*Lats2*^-/-^-EKO, by crossing *Cdh5-cre/ERT2* mice with *Lats1*^fl/fl^*Lats2*^fl/fl^ mice on a C57BL/6 background. Non-knockout littermate controls (NLC) were used for comparison. Deletion efficiency was validated one-week post-induction. Western blotting of primarily lung endothelial cells (MLECs) confirmed loss of LATS1/2 protein (**Fig. 1A**).

**Figure 1.**
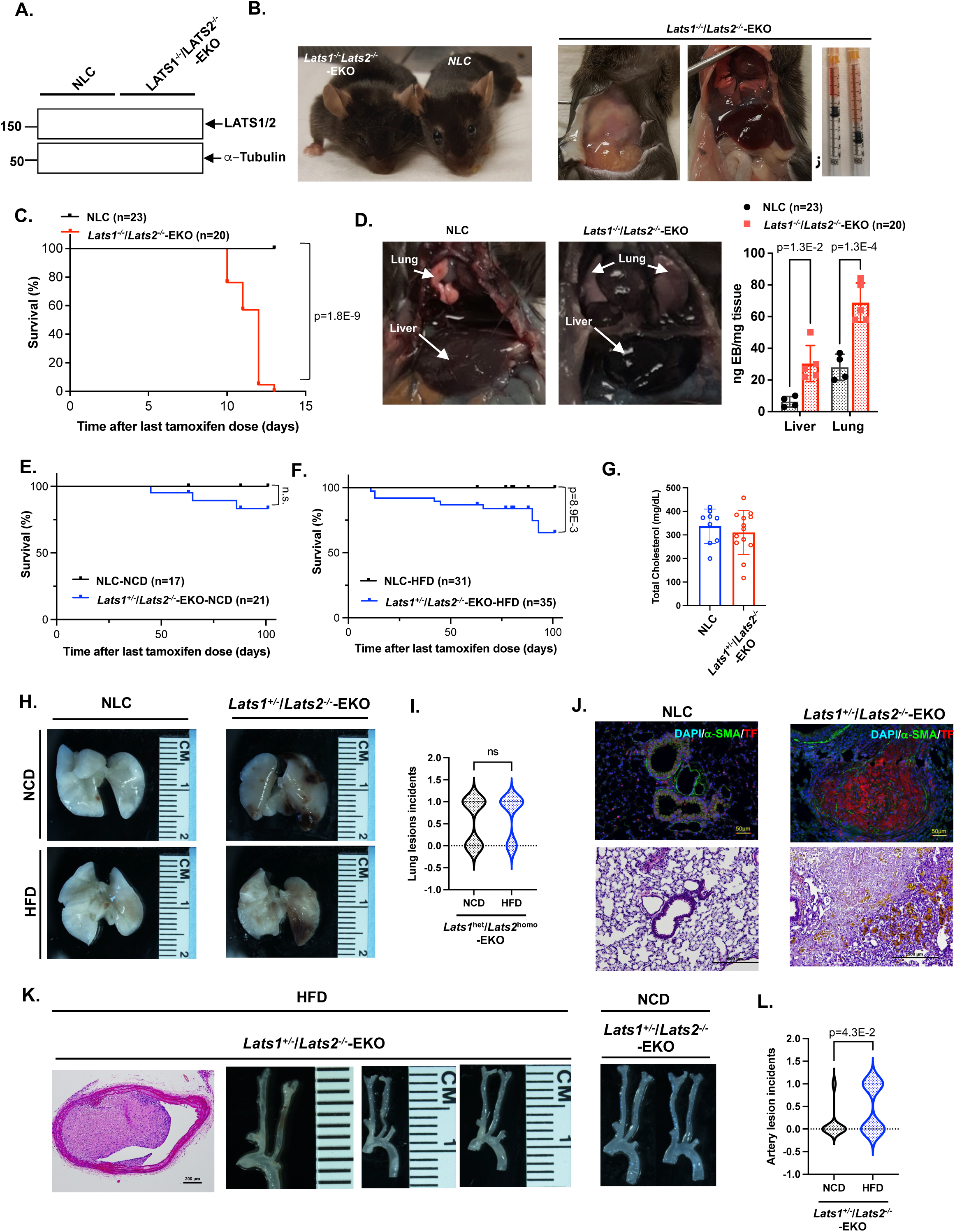
Endothelial LATS1/2 are essential for vascular barrier integrity and survival. **(A)** Mice were administered tamoxifen (75 mg/kg, intraperitoneally for five consecutive days) at six weeks of age. After 8-days of the final tamoxife injection, mouse lung endothelial cells (MLECs) were isolated for Immunoblot analysis of LATS1/2 protein expression. Samples were obtained from tamoxifen-induced *Lats1*^-/-^/*Lats2*^-/-^- endothelial-specific knockout (EKO) mice and non-transgeneic littermate control (NLC). **(B)** Representative gross images of *Lats1*^-/-^/*Lats2*^-/-^-EKO mice 12 days post-tamoxifen induction, demonstrating facial edema, pleural effusion, and hemorrhagic ascites. Cre recombination efficiency is validated in parallel. **(C)** Kaplan-Meier survival curve showing that all *Lats1*^-/-^/*Lats2*^-/-^-EKO mice (n = 20) succumbed within 15 days post-tamoxifen, whereas all NLC mice (n = 23) survived. **(D)** Gross necropsy of liver and lung tissues reveals vascular congestion and edema in *Lats1*^-/-^/*Lats2*^-/-^-EKO mice, confirmed by Evans Blue dye extravasation quantification (n = 4–5/group). **(E–F)** Survival curves of *Lats1*^+/−^/*Lats2*^-/-^-EKO and NLC mice under NCD (E) and hyperlipidemic stress (AAV-PCSK9 + HFD) (F), indicating reduced survival in *Lats1*^+/−^/*Lats2*^-/-^- EKO mice under HFD. **(G)** Serum cholesterol levels in *Lats1*^+/−^/*Lats2*^-/-^-EKO versus NLC mice on HFD, confirming no significant differences (n = 9-13/group; unpaired t-test). **(H)** Representative gross images of lungs from NLC and *Lats1*^+/−^/*Lats2*^-/-^-EKO mice under NCD and HFD. Thrombus-like consolidations are apparent in EKO lungs irrespective of diet. **(I)** Quantification of pulmonary thrombus incidence across diets reveals diet-independent lesion frequency in *Lats1*^+/−^/*Lats2*^-/-^-EKO mice (ns, not significant). (NCD, total n=19, 1: n=9, 0: n=7, HFD, total n=13, 1: n=9, 0: n=4) **(J)** Immunofluorescence and histological staining of lung sections from NLC and *Lats1*^+/−^/*Lats2*^-/-^-EKO mice. α-SMA and DAPI staining (top) show vascular remodeling; H&E (bottom) shows lung infarction resulting from pulmonary thrombosis and disrupted architecture. **(K)** Cross-sectional and whole-mount images of carotid arteries from *Lats1*^+/−^/*Lats2*^-/-^-EKO mice reveal luminal narrowing under both HFD (left) and NCD (right). **(L)** Quantification of luminal cross-sectional area confirms vessel narrowing in *Lats1*^+/−^/*Lats2*^-/-^- EKO mice under NCD. (NCD, total n=17, 1: n=2, 0: n=15, HFD, total n=13. 1: n=6, 0: n=7). Statistical information—including the sample size (n) for each group, results of normality tests, statistical methods applied, and corresponding P values—is also summarized in Table S1. All data are expressed as mean±SD.

By day 12 post-induction, *Lats1*^-/-^/*Lats2*^-/-^-EKO mice developed severe systemic illness marked by facial edema, pleural effusion, and hemorrhagic ascites (**Fig. 1B, left** and **middle**). Gross necropsy revealed fluid accumulation in thoracic and abdominal cavities. The peritoneal fluid appeared serosanguinous, consistent with systemic vascular leakage-a phenotype absents in control animals (**Fig. 1B, right).** All *Lats1*^-/-^/*Lats2*^-/-^-EKO mice (n=20) succumbed within 15 days post-induction, while all NLC mice (n=23) survived (**Fig. 1C**). Evans Blue extravasation assays showed increased vascular permeability in the lungs and liver of *Lats1*^homo^/*Lats2*^homo^-EKO ten days post-tamoxifen (**Fig. 1D**).

Serum markers of liver and renal function remained within physiological ranges (**Fig. S1A**), and no significant cardiac dysfunction was detected within 6 days following tamoxifen injection (**Fig. S1B**). These findings suggest that the systemic edema observed in these mice is unlikely to be attributable to hepatic, renal, or cardiac dysfunction—at least within the first 6 days after tamoxifen induction.

### Pulmonary thrombosis and arterial lesions in *Lats1*<u>^het(+/−)^</u>/*Lats2*^homo(-/-)^-EKO mice under hypercholesterolemia

Since *Lats1*^-/-^/*Lats2*^-/-^-EKO mice do not survive, we evaluated the phenotype of *Lats1***<u>^het(+/−)^</u>**/*Lats2*^homo(-/-)^-EKO mice. After tamoxifen administration, we did not observe systemic edema in these mice, and their survival rate was similar to NLC mice (**Fig. 1E**). Given the potential role of endothelial LATS1/2 in maintaining vascular barrier integrity, we investigated whether hypercholesterolemia exacerbates disease progression. To induce atherogenic stress, we injected *Lats1***^+/−^**/ *Lats2*^-/-^-EKO with AAV8-PCSK9 and started a high-fat diet (HFD) as we performed previously^20^. We found that *Lats1***^+/−^** / *Lats2* ^-/-^ -EKO mice under hypercholesterolemic conditions exhibited significantly reduced survival compared to WT mice (**Fig. 1F**). Serum cholesterol levels were comparable between genotypes (**Fig. 1G**). Additionally, there was no significant difference in body weight between *Lats1***<u>^+/−^</u>**/ *Lats2*^-/-^-EKO mice and NLC (**Fig. S1C**).

Notably, *Lats1***^+/−^**/*Lats2*^-/-^-EKO mice developed spontaneous pulmonary thrombi under both normal chow diet (NCD) and high-fat diet (HFD), evident as opaque, consolidated regions in the lungs (**Fig. 1H**). The frequency of these lesions was similar across diets (**Fig. 1I**), indicating that partial endothelial depletion of LATS1/2 alone is sufficient to induce thrombosis, independent of dietary lipid levels. Immunostaining for α-smooth muscle actin (α-SMA) and tissue factor (TF), a key pro-coagulant protein, revealed thrombus-like occlusions within pulmonary vessels of *Lats1***^+/−^** /*Lats2*^-/-^-EKO mice, regardless of dietary condition. Corresponding hematoxylin and eosin (H&E) staining confirmed luminal occlusion, lung infarction, and disrupted vascular architecture, consistent with thrombotic remodeling and hemosiderin deposition (**Fig. 1J**).

Arterial lesions were observed in the aorta and carotid arteries of mice on both NCD and HFD with AAV-PCSK9 injection in *Lats1***<u>^+/−^</u>**/*Lats2*^-/-^-EKO mice. We did not find significant differences of total cholesterol level, and body weight (BW) between WT and *Lats1***^+/−^**/*Lats2*^-/-^- EKO mice (**Fig. 1G, S1C**). However, the prevalence of arterial lesions was significantly higher in mice on the HFD compared to those on the NCD (**Fig. 1K, L**). These arterial lesions may contribute to the worsened survival observed in *Lats1***^+/−^**/*Lats2*^-/-^-EKO mice under HFD conditions (**Fig.1F**).

### Distinct atherothrombosis formation with enhanced neoangiogenesis and immune cell infiltration in carotid arteries subjected to d-flow in *Lats1*^+/−^/*Lats2*^-/-^-EKO mice

In *Lats1***^+/−^**/*Lats2*^-/-^-EKO mice, spontaneous arterial lesions were found in both aorta and carotid arteries, albeit inconsistently of the lesion locations, making direct comparison with NLC mice challenging due to the variable lesion locations (**Fig. 1K**). Consequently, to establish a controlled comparison, we executed a partial left carotid ligation (PLCL) on both *Lats1***^+/−^**/*Lats2*^-/-^-EKO mice and NLC mice (**Fig. 2A**). Fig. 2B-D illustrates that the post-PLCL lesion size in *Lats1***^+/−^**/*Lats2*^-/-^-EKO mice after 2 weeks of PLCL surgery was significantly larger compared to that in NLC mice, a finding consistent across both sexes.

**Figure 2.**
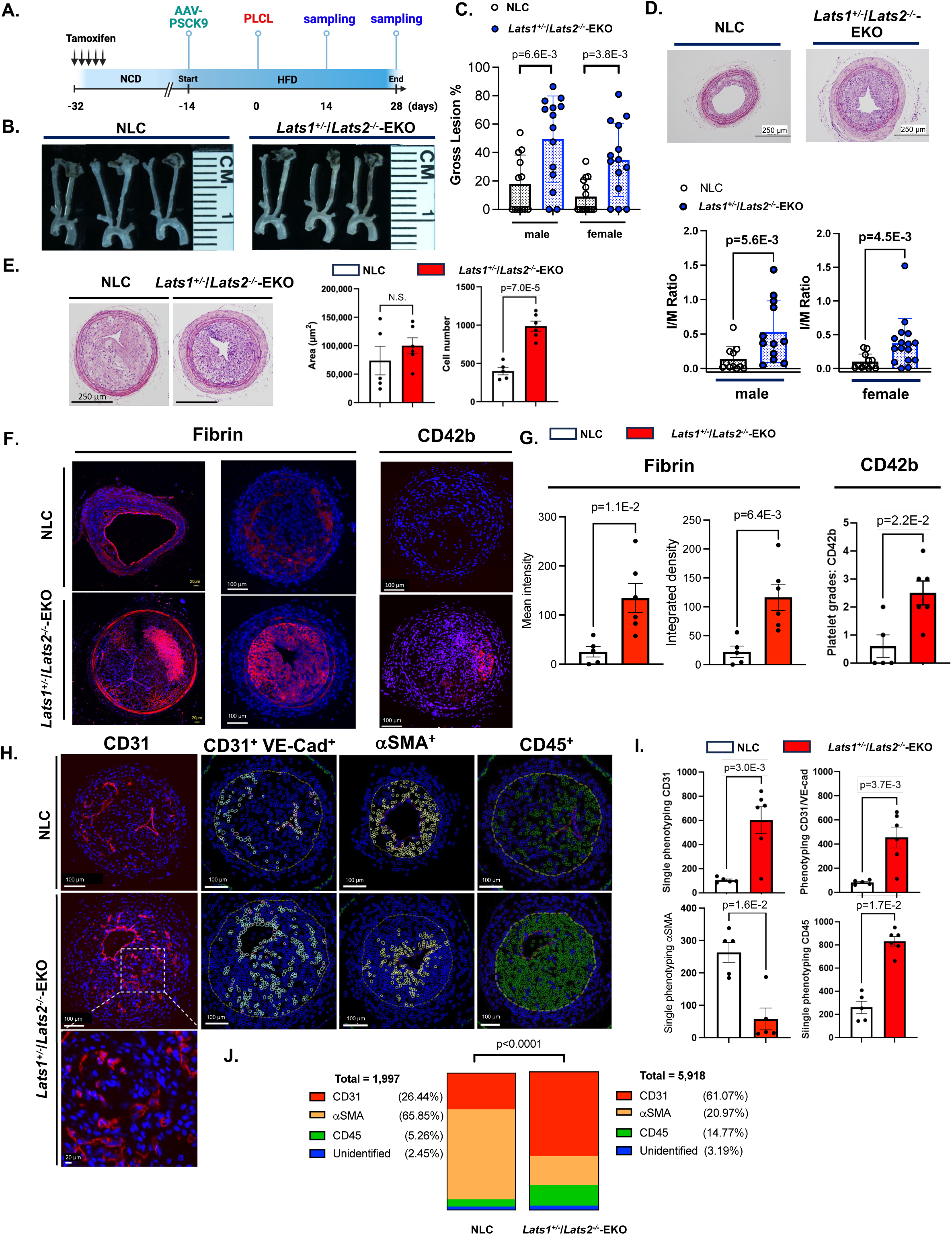
Plaques in *Lats1*^+/−^/*Lats2*^-/-^-EKO mice exhibit atherothrombosis and enhanced neo-angiogenesis. **(A)** Experimental design for PLCL. Tamoxifen was administrated for five consecutive days, and experiments were initiated 15 days after the final injection. NLC and *Lats1*^+/−^/*Lats2*^-/-^-EKO mice were administered AAV-PCSK9 followed by HFD and PLCL to induce d-flow in the left carotid artery (LCA), with the right carotid artery (RCA) serving as a static flow control **(B-C)** Gross plaque area quantification from whole-mount carotid arteries in male and female NLC and *Lats1*^+/−^/*Lats2*^-/-^-EKO mice, 2 weeks post-PLCL. *Lats1*^+/−^/*Lats2*^-/-^-EKO mice show increased plaque burden under d-flow. **(D)** Representative H&E-stained cross-sections of LCA and RCA from NLC and *Lats1*^+/−^/*Lats2*^-/-^-EKO mice harvested 2 weeks post-PLCL. Bottom: Plaque burden was quantified as intima plus media divided by media (I/M ratio) separately in male and female mice. Scale bars, 250 µm. **(E)** To compare plaque phenotype between groups independent of plaque size, *Lats1*^+/−^/*Lats2*^-/-^- EKO mice were harvested 2 weeks post-PLCL and NLC mice at 4 weeks, resulting in size-matched lesions. Although intimal plaque area were similar, total cell number within plaques was increased in *Lats1*^+/−^/*Lats2*^-/-^-EKO mice. **(F, G)** Representative immunofluorescence staining of fibrin and CD42b in plaques from NLC and *Lats1*^+/−^/*Lats2*^-/-^-EKO mice. Increased deposition of fibrin (G: mean intensity and integrated density as described in the Methods) and platelet marker CD42b is observed in *Lats1*^+/−^/*Lats2*^-/-^- EKO mice lesions. A grading system was used to evaluate CD42b immunofluorescence staining in the plaques, as described in the Methods (G). Data are presented as mean ± SD, n=5-6. **(H)** COMET™ imaging was followed by automated segmentation and quantification of CD31⁺ ECs, VE-Cadherin⁺ double-positive ECs, αSMA⁺ ⁺ vascular smooth muscle cells (VSMCs),, and CD45⁺ leukocytes within in LCA plaques. **(I)** Quantification of individual cell populations (CD31⁺, CD31⁺/VE-Cadherin⁺, αSMA⁺, CD45⁺) in plaques from NLC and *Lats1*^+/−^/*Lats2*^-/-^-EKO plaques. EC and leukocyte counts were significantly elevated in *Lats1*^+/−^/*Lats2*^-/-^-EKO lesions, whereas VSMCs were reduced (n = 5– 6/group; mean ± SD). **(J)** Stacked bar plots showing relative proportions of CD31⁺ ECs, αSMA⁺ VSMCs, and CD45⁺ leukocytes among all segmented plaque cells in each genotype. *Lats1*^+/−^/*Lats2*^-/-^-EKO plaques are enriched for ECs and leukocytes, with reduced VSMC content. Statistical information—including the sample size (n) for each group, results of normality tests, statistical methods applied, and corresponding P values—is also summarized in Table S1.

To accurately assess and compare the characteristics of arterial lesions in *Lats1***^+/−^**/*Lats2*^-/-^-EKO and NLC mice, we matched lesion sizes in NLC mice observed at 4 weeks post-PLCL with those in *Lats1***^+/−^**/*Lats2*^-/-^-EKO mice at 2 weeks post-PLCL. This approach ensures that any observed differences in lesion characteristics are attributed to the LATS1/2 deletion rather than to differences in lesion size (**Fig. 2E**). As expected, we did not find any differences in lesion sizes between *Lats1***^+/−^**/*Lats2*^-/-^-EKO mice (2 weeks after PLCL) and NLC (4 weeks after PLCL) (**Fig. 2E)**. Prompted by the pronounced upsurge in pulmonary thrombotic occurrences in the *Lats1***^+/−^**/*Lats2*^-/-^-EKO mice, we extended our scrutiny to the thrombotic elements within arterial lesions by using IMC/COMET^TM^ technology. This examination revealed a substantial elevation in the expression of fibrin, as well as CD42b, a platelet marker, within the carotid lesions of the *Lats1***<u>^+/−^</u>**/*Lats2*^-/-^-EKO mice, relative to the NLC group (**Fig. 2F, G**). Subsequent to our examination of the cellular makeup within both types of lesions, we observed a marked increase in the prevalence of CD31^+^ endothelial cells and CD45^+^ leukocytes, alongside a notable reduction of α- SMA^+^ cells in the *Lats1*^+/−^/*Lats2*^-/-^-EKO mice when juxtaposed with their NLC counterparts (**Fig. 2H-J**). These findings indicate that the lesions in *Lats1***^+/−^**/*Lats2*^-/-^-EKO mice are characterized by an increased abundance of hematopoietic and thrombotic components, strong infiltration of leukocytes, enhanced neoangiogenesis, and a decreased presence of myofibroblasts and vascular smooth muscle cells, collectively reflecting features typical of atherothrombotic lesions.

### CD38 links senescence and proliferation in LATS1/2-depleted endothelial cells

In the ECs marked by CD31^+^ within *Lats1*^+/−^/*Lats2*^-/-^-EKO mice, a downregulation of LATS1 and upregulation of YAP expression were noted (**Fig. S2A**), corroborating the in vivo depletion of endothelial LATS1. We also found the expression of tissue factor (TF) was significantly upregulated in ECs from *Lats1*^+/−^/*Lats2*^-/-^-EKO mice (**Fig. S2B**). Additionally, there was a significant increase in senescence-associated secretory (SASP)-related factors, including inflammatory markers such as IL-6, CCR7, CD44 (**Fig. S2C**), and the senescence marker p53 (**Fig. S2E**), along with the proliferation marker Ki67 and SPP1 (**Fig. S2F**), were detected. In contrast, vimentin expression was not increased, suggesting that endothelial-to-mesenchymal transition (EndoMT) is unlikely to contribute significantly in *Lats1*^+/−^/*Lats2*^-/-^-EKO mice (**Fig. S2D**).

Because both the senescence marker p53 and the proliferation marker Ki67 were elevated in ECs from *Lats1*^+/−^/*Lats2*^-/-^-EKO mice, we investigated whether there was a relationship between these two markers. This analysis aimed to explore the possibility of senescence-associated stemness (SAS), a concept we previously described in the context of plaque formation in macrophages^21^. However, the correlation between p53 and Ki67 expression was weak in both NLC mice (R² = 0.039) and *Lats1*^+/−^/*Lats2*^-/-^-EKO mice (R² = 0.0014) (**Fig. S3A**). This finding suggests that p53 and Ki67 are not directly linked in this setting.

In addition to p53, we identified other markers associated with SASP using imaging mass cytometry (IMC) and COMET analysis. Among these, we focused on CD38, an enzyme involved in NAD⁺ metabolism that has been implicated in both cellular senescence and proliferation^22,23^. Given this dual role, we investigated the relationship between CD38 and the proliferation marker Ki67. We observed a significant upregulation of CD38 expression in ECs within atherosclerotic plaques from Lats1⁺/⁻/Lats2⁻/⁻-EKO mice (**Fig. 3A**). To further elucidate the role of CD38 in regulating cell proliferation, we performed single-cell co-expression analysis of CD38 and Ki67 using IMC/COMET, as described in the Methods section (**Fig. 3B–E**). By computing the log ratio of Ki67 to CD38 expression, we observed a trimodal distribution (**Fig. 3C**). In NLC ECs, two distinct cell populations were identified based on CD38 and Ki67 expression: Group 1, with high CD38 and low Ki67, and Group 3, with low CD38 and high Ki67 (**Fig. 3B**). These mutually exclusive patterns are consistent with the established view of CD38 as a marker of cellular senescence, which is typically linked to reduced proliferative capacity. In contrast, LATS1/2-deficient ECs from *Lats1*^+/−^/*Lats2*^−/−^-EKO mice exhibited a third population—Group 2—defined by high expression of both CD38 and Ki67. This co-expression showed a linear correlation between the two markers, suggesting a breakdown in the typical inverse relationship between senescence and proliferation. Notably, the proportion of Group 2 cells was significantly higher in LATS1/2-deficient ECs compared to non-lesion controls **(Fig. 3D, E)**. These findings indicate that CD38 may acquire a pro-proliferative role in the absence of LATS1/2, allowing cells to adopt a dual phenotype of senescence and proliferation, deviating from its conventional function in promoting cell cycle arrest.

**Figure 3.**
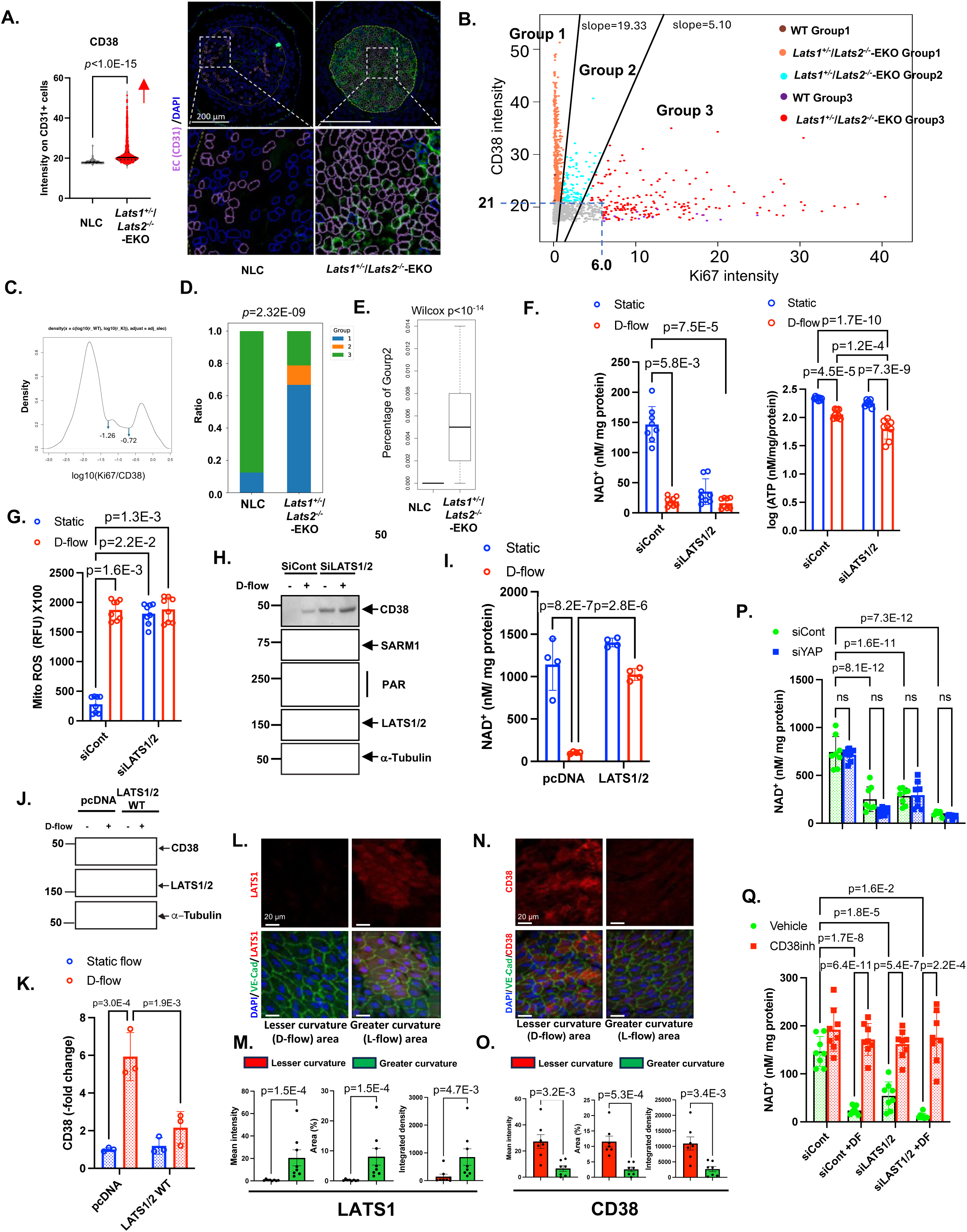
Upregulation of the senescence-associated molecule endothelial CD38 in plaques of *Lats1*<u>^+/−^</u>/*Lats2*^-/-^-EKO mice drives NAD^+^ depletion in response to d-flow exposure. **(A)** CD38 expression levels in CD31^+^ ECs were significantly higher in *Lats1***<u>^+/−^</u>**/*Lats2*^-/-^-EKO mice compared to NLC mice. **(B)** Single-cell analysis showing the expression levels of CD38 and Ki67 in ECs in vivo. **(C)** Logarithmic ratio of Ki67 to CD38 expression in selected cells, calculated as log10(Ki67 expression/CD38 expression). The cutoff values for the ratio are set at log10(−1.26) and log10(−0.72). **(D)** Stacked bar charts illustrating the proportions of Group 1–3 ECs (defined in panel B) within plaques of NLC and *Lats1***<u>^+/−^</u>**/*Lats2*^-/-^-EKO mice. **(E)** Percentage distribution of ECs in Groups 1–3 within plaques of NLC and *Lats1***<u>^+/−^</u>**/*Lats2*^-/-^- EKO mice. Resampling was performed 1,000 times, selecting 500 cells from each group in each round to calculate the percentage of cells in Group 1, 2, or 3. The percentage of cells in Group 2 was calculated as (# of Group 2 cells / # of total cells). Statistical significance was assessed using the Wilcoxon test, as described in the Methods. **(F-H)** HUVECs transfected with siCont or siLATS1/2 were exposed to d-flow for 24 hours. Levels of NAD^+^ and ATP levels (F), as well as mtROS (G) were measured. Protein expression was analyzed by Western blot using specific antibodies (H). **(I-K)** HUVECs transfected with plasmids expressing LATS1 and LATS2 or a control vector (pcDNA) were exposed to d-flow for 24 hours. NAD^+^ levels measured(I) and protein expression of CD38, LATS1, and LATS2 (J) were measured. **(K)** The graphs represent densitometric data from three independent blots. **(L-O)** En face preparations of the aortic arch from 10-week-old WT C57BL/6 mice stained with anti-VE-Cad (an EC marker) and anti-LATS1 (L, M), anti-CD38 (N, O). Scale bars = 20 µm. Quantification of LATS1 and CD38 expression is presented as mean intensity, area (%), and integrated density (sum of pixel intensity) as described in Methods in bar graphs; n = 7-8 mice. **(P, Q)** HUVECs transfected with siCont or siYAP (P) or pre-treated with the CD38 inhibitor 78c (1 µM) (Q) were exposed to d-flow for 24 hours. NAD^+^ levels were measured. Statistical information—including the sample size (n) for each group, results of normality tests, statistical methods applied, and corresponding P values—is also summarized in Table S1.

### D-flow induces CD38 expression via the inhibition of LATS1/2 expression, and subsequently induces NAD^+^/ATP^+^ depletion and mitochondrial reactive oxygen species (ROS) production

We observed that LATS1/2 depletion selectively enhances d-flow-induced CD38 expression, as well as NAD^+^/ATP depletion and mitochondrial ROS (mtROS) induction (**Fig. 3F, G**). Notably, this regulation does not affect the expression or activation of other NAD⁺ hydrolases, such as SARM1 and poly(ADP-ribose) polymerase (PARP) (Fig. 3H, S3C). Conversely, LATS1/2 overexpression mitigates d-flow-induced NAD^+^ depletion (**Fig. 3I**) and suppresses CD38 expression (Fig. 3J–K). In vivo, we observed a concurrent downregulation of LATS1 and upregulation of CD38 expression in the lesser curvature (disturbed-flow region) of the wild type mouse aortic arch, compared to the greater curvature (laminar-flow region) (**Fig. 3L–O**).

To further investigate the impact of CD38 induction following LATS1/2 depletion, we examined its role in d-flow-mediated NAD⁺ depletion and compared it with YAP activation. While CD38 inhibition fully restored NAD^+^ levels depleted by d-flow and LATS1/2 loss, YAP reduction had no significant effect (**Fig. 3P, Q**), indicating that CD38, rather than YAP, is a primary regulator of NAD^+^ depletion in this setting.

### Inhibition of CD38 Activity, not YAP depletion, reduces all senescence markers induced by d-flow through Lamin A reduction

Although the depletion of YAP inhibited the increase in p21, ICAM1, and TF expression, as well as the number of Ki67 positive and apoptotic cells following d-flow in ECs with depleted LATS1/2, no significant differences were observed in other tested SASP events (**Fig. 4A-D, S4A**). Conversely, administering a specific CD38 inhibitor significantly mitigated the effects of d-flow on all investigated SASP, proliferation, and apoptosis markers, as well as YAP transcription TEAD activity (**Fig. 4E-I, S4B**). These findings suggest that, unlike CD38, YAP cannot regulate the entire set of senescence-related events, including NAD^+^ depletion. However, both CD38 and YAP significantly impact EC inflammation and proliferation after d-flow. Suppressing CD38 leads to an increase in LATS1/2 expression, indicating a positive feedback mechanism involving CD38 (**Fig. 4E, S4B**). This evidence underscores the primary role of CD38 in regulating both upstream and downstream events of LATS1/2 degradation, a capability that YAP does not possess.

**Figure 4.**
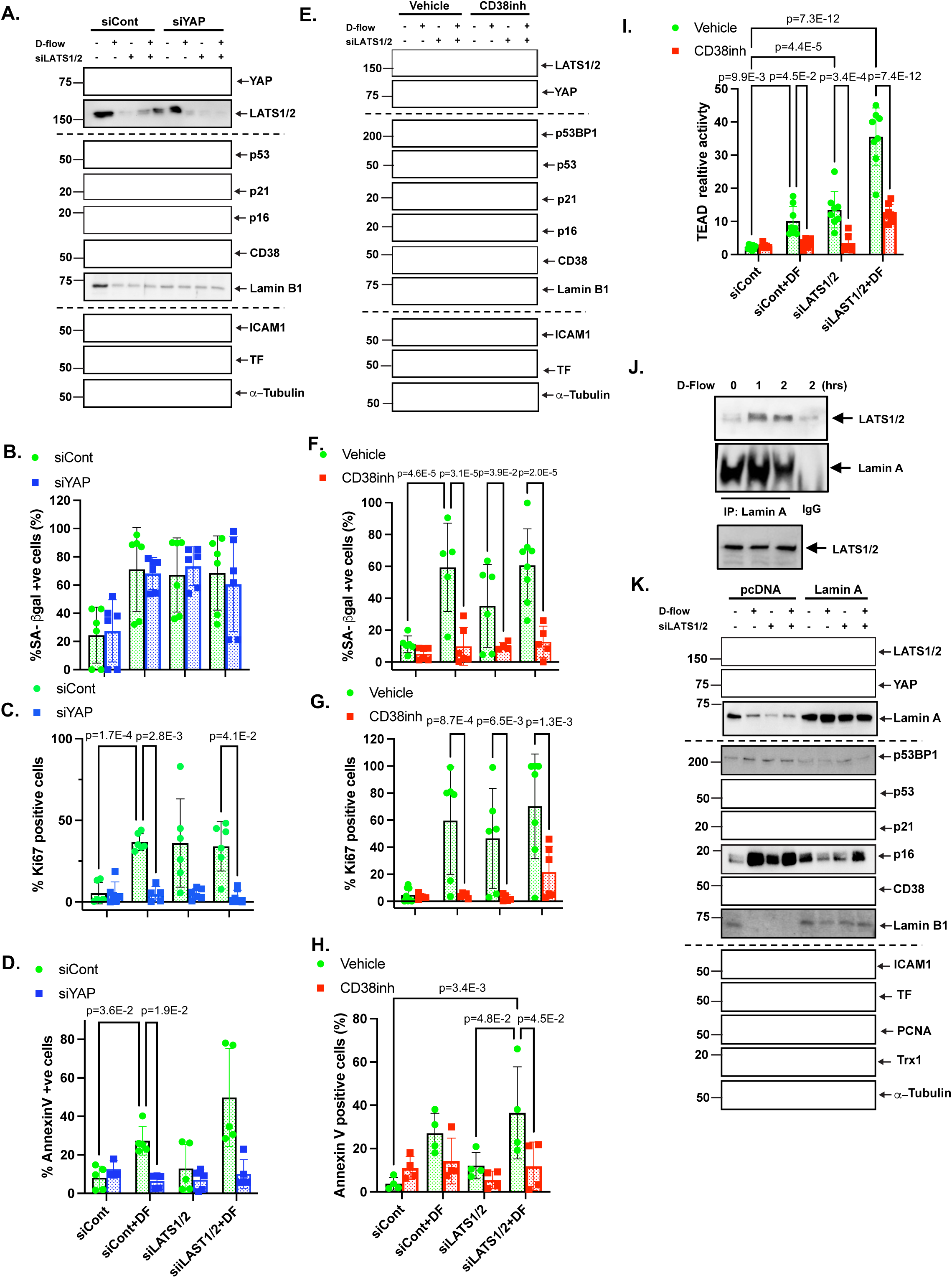
Under d-flow conditions, CD38 and YAP function through distinct mechanisms, with CD38 simultaneously inducing senescence, inflammation, apoptosis, and proliferation via a pathway regulated by Lamin A in association with LATS1/2. **(A-D)** HUVECs were transfected with siCont, siYAP, siLATS1/2, or a combination of siYAP and siLATS1/2, and then exposed to d-flow or maintained under static conditions for 24 hours. Protein expression was analyzed by Western blot using specific antibodies (A). The percentages of SA-β-gal^+^ cells (Senescence) (B), Ki67^+^ cells (Proliferation) (C), and annexin V^+^ cells (apoptosis) (D) were measured by flow cytometry. **(E-H)** HUVECs transfected with siCont or siLATS1/2, followed by treatment with or without pre-treatment with the CD38 inhibitor 78c (1 µM), were exposed to d-flow or maintained under static conditions. Protein expression was analyzed by Western blot using specific antibodies (E). The percentages of SA-β-gal^+^ cells (F), Ki67^+^ cells (G), and annexin V^+^ cells (H) were detected by flow cytometry. **(I)** HUVECs transfected with siCont or siLATS1/2 and a plasmid containing a TEAD activity reporter gene were exposed to d-flow or maintained under static conditions for 8 hours. TEAD activity was measured using a luciferase assay. **(J)** HUVECs exposed to d-flow for the indicated durations were subjected to immunoprecipitation with anti-Lamin A or IgG control antibodies, followed by Western blotting with anti-LATS1/2 antibodies to detect potential binding. To confirm equal immunoprecipitation specificity, the blot was subsequently stripped and reprobed with anti-lamin A. Immunoprecipitation with control IgG served as a negative control to validate the specificity of the interaction **(K)** HUVECs transfected with siCont or siLATS1/2, along with a plasmid expressing Lamin A or an empty vector, were exposed to d-flow, and protein expression was analyzed by Western blot using specific antibodies. Quantification data for panels A, E, J, and K are provided in the corresponding figure panels in Fig. S4. Data are presented as mean ± SD. Statistical tests, sample sizes, and detailed results for all panels are summarized in Table S1.

The reduction of Lamin A expression with aging can compromise the structural integrity of the nucleus, leading to cellular senescence and tissue dysfunction, thereby contributing to the aging phenotype^24^. The association between LATS1/2 and Lamin A has been documented^25^, with this association being upregulated after 1 hour of d-flow (**Fig. 4J, S4C**). However, after 24 hours of d-flow and LATS1/2 depletion, the expression of both was significantly reduced (**Fig. 4K, S4D**). While the binding of Lamin A to the CD38 promoter region has been observed^26^, its functional role remains unclear. Overexpression of Lamin A was found to inhibit the induction of CD38 and subsequent SASP caused by d-flow in ECs depleted of LATS1/2 (**Fig. 4K, S4D**). This supports the role of Lamin A in CD38-mediated SASP induction, which appears to occur independently of YAP activation. These findings suggest that CD38 can drive divergent cellular outcomes—senescence and proliferation—depending on the functional status of LATS1/2. In the presence of functional LATS1/2, CD38 is associated with a senescence program, potentially mediated through Lamin A and the SASP. However, in the absence of LATS1/2, CD38 may simultaneously support proliferative signaling, indicating that proliferation can occur even in the context of senescence (**Fig. 5L**).

**Figure 5.**
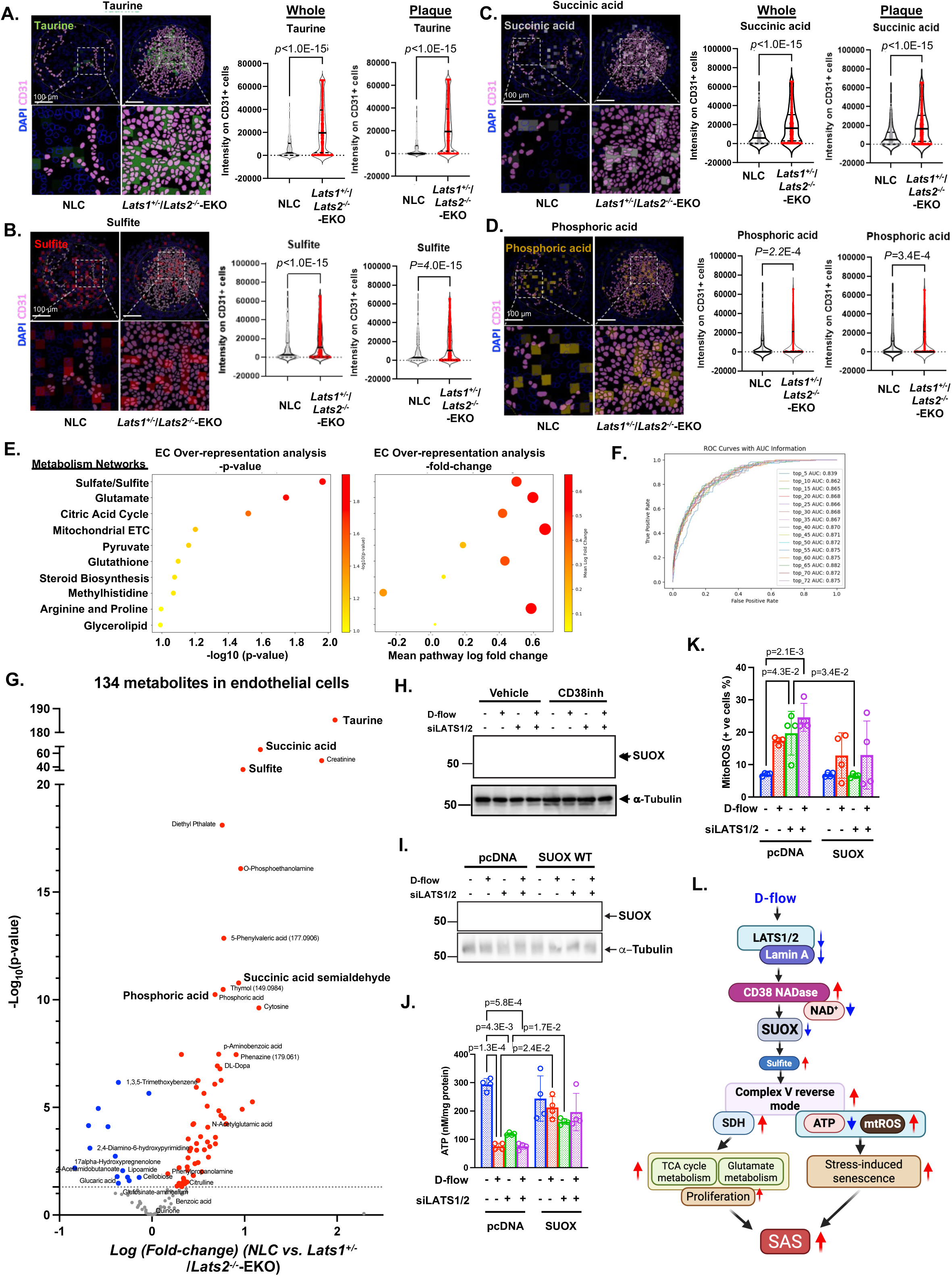
Spatial metabolomics analysis revealed increased sulfate/sulfite metabolism networks in ECs of *Lats1*<u>^+/−^</u>/*Lats2*^-/-^-EKO mice compared to NLC, highlighting the role of SUOX in d-flow-induced ATP reduction, decreased ATP synthase activity, and elevated mtROS levels. **(A-E)** By integrating IMC/COMET™ and spatial metabolomics data, single-cell metabolite profiles were obtained for ECs. Analysis revealed that sulfate/sulfite metabolism (Taurine (A), sulfite (B)), glutamate metabolism (succinic acid (C), phosphoric acid (D)), and the citric acid (TCA) cycle metabolism networks were disproportionately represented and significantly upregulated in ECs of *Lats1*<u>^+/−^</u>/*Lats2*^-/-^-EKO mice (E) compared to NLC mice. The “whole” region encompasses the ECs lining the vessel surface, while the “plaque” region specifically excludes the endothelial surface. **(F)** Receiver operating characteristic (ROC) curve demonstrating the predictive ability of EC-specific metabolite data to distinguish *Lats1*<u>^+/−^</u>/*Lats2*^-/-^-EKO mice from NLC mice. **(G)** Volcano plot illustrating the 134 metabolites detected in ECs from spatial metabolomics analysis of plaques from *Lats1*<u>^+/−^</u>/*Lats2*^-/-^-EKO mice compared to NLC mice. **(H)** HUVECs transfected with siCont or siLATS1/2, with pre-treatment using the CD38 inhibitor 78c (1 µM) or vehicle, were exposed to d-flow, and protein expression was analyzed by Western blot. **(I-K)** HUVECs transfected with siCont or siLATS1/2, along with a plasmid expressing WT SUOX or an empty vector (pcDNA), were exposed to d-flow for 24 hours. Protein expression was analyzed by Western blot (I), and measurements of ATP levels (J), and mtROS levels (K) were performed as described in the Methods. Statistical tests, sample sizes, and detailed results for all panels are summarized in Table S1. (**L**) **Graphical abstract:** Lamin A binds directly to the CD38 promoter, repressing its transcription. Under physiological conditions, LATS1/2 and Lamin A work together to suppress CD38 expression. LATS1/2 can interact with Lamin A, and their combined action reinforces CD38 repression. Therefore, when d-flow downregulates both LATS1/2 and Lamin A, this dual repression is lifted, leading to increased CD38 expression. CD38, in turn, modulates SUOX expression, either directly or through feedback involving LATS1/2. SUOX controls Complex V (CV) reverse mode, which increases mitochondrial ROS production and ATP depletion, while CD38-driven NAD⁺ consumption further exacerbates metabolic stress. In response, there is a compensatory upregulation of succinate dehydrogenase (SDH), enhancing TCA cycle activity and glutamate metabolism, thereby generating biosynthetic precursors for protein and nucleic acid synthesis. This supports a paradoxical state in which stress-induced senescence coexists with proliferative endothelial behavior. Together, these coordinated events define a non-canonical senescence program—senescence-associated sprouting (SAS)—driven by the LATS1/2–CD38–SUOX axis.

### Spatial metabolomics findings indicate SUOX deficiency leads to upregulation of Sulfate/Sulfite, Glutamate, and TCA Cycle metabolism networks in *Lats1*^het^/*Lats2*^homo^-EKO Mice

Building on the observed NAD^+^/ATP depletion, we next sought to identify the metabolic pathways involved in the CD38–LATS1/2 regulatory axis. To this end, we integrated spatial metabolomics with IMC/COMET data using the VISIOPHARM analysis platform. This approach enabled the identification of 134 distinct metabolites in ECs. We performed an overrepresentation analysis on single CD31^+^ cell-based spatial metabolomics data. We found that the metabolite networks of sulfate/sulfite, glutamate, and the TCA cycle in ECs were significantly enhanced in the plaques of *Lats1*^+/−^/*Lats2*^-/-^-EKO mice compared to NLC mice (**Fig. 5A-E**). Our predictive model showed that these metabolites in ECs could predict the phenotype of *Lats1*^+/−^/*Lats2*^-/-^-EKO mice with an area under the curve (AUC) of 0.882 (**Fig. S5A**). SHAP (SHapley Additive exPlanations) analysis helps interpret complex models by assigning each feature an importance value for a particular prediction^27^, which also revealed that the top three metabolite networks mentioned above had a stronger predictive power for the phenotype of *Lats1*^+/−^/*Lats2*^-/-^-EKO mice compared to other molecules detected by IMC/COMET (**Fig. S5B**). This finding underscores the crucial role these three metabolite networks play in determining the unique phenotype of ECs in the plaques from *Lats1*^+/−^/*Lats2*^-/-^-EKO mice.

Given the marked increases in both taurine and sulfite levels (**Fig. 5G**)—metabolic changes consistent with SUOX deficiency^28,29^—we examined SUOX expression. We found that SUOX expression was significantly reduced in ECs exposed to d-flow and upon LATS1/2 depletion. Notably, this reduction was attenuated by treatment with a CD38 inhibitor (**Fig. 5H, S5C**). Since elevated sulfite resulting from SUOX deficiency can cause sulfite toxicity through thiol modification of the F0/F1-ATP synthase complex—ultimately impairing ATP production^30^—we next assessed ATP levels. We observed that d-flow and/or LATS1/2 depletion significantly reduced ATP levels and increased mtROS. These effects were reversed by SUOX overexpression (**Fig. 5 I-K**), supporting a critical role for SUOX in maintaining ATP production and limiting mtROS generation under conditions of d-flow and LATS1/2 depletion (**Fig. 5L**).

Next, we assessed metabolic flux using ex vivo [¹³C]glucose and [¹³C]glutamine tracing in ECs isolated from the left (LCA, d-flow) and right (RCA, l-flow) carotid arteries seven days after PLCL. After a 24-hour recovery, ECs were incubated with tracers for 24 hours, followed by LC- MS analysis (**Fig. 6A**). Glutamine tracing showed similar ¹³C incorporation into glutamate and α- ketoglutarate in both groups, but significantly reduced labeling of downstream TCA metabolites in LCA ECs, indicating impaired OGDH activity (**Fig. 6B**). Glucose tracing revealed comparable glycolytic flux, but reduced TCA cycle labeling in LCA ECs (**Fig. 6C**), linked to decreased PDH expression and activity. This reduction was further exacerbated by LATS1/2 depletion and was CD38 dependent (**Fig. 6D, E**). Overall, disturbed flow impairs OGDH and PDH activity, disrupting TCA cycle metabolism in ECs.

**Figure 6.**
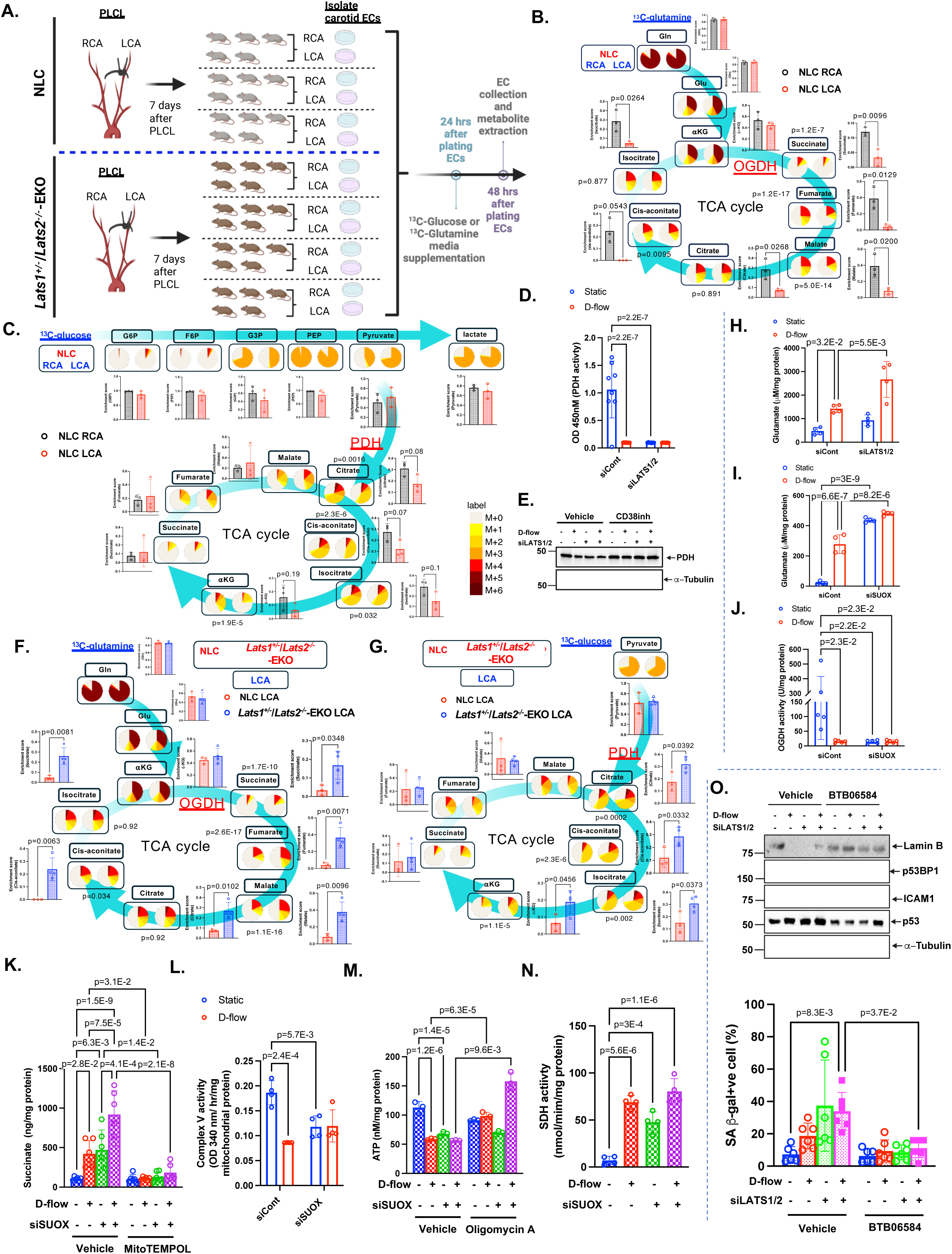
D-flow suppresses oxoglutarate dehydrogenase (OGDH) and pyruvate dehydrogenase (PDH) activity ex vivo, but SUOX deficiency in LATS1/2 EKO induces compensatory effects through activating complex V reverse mode, enhancing glutamate metabolism and TCA cycle activation. (**A, B, C, F, G**) To investigate metabolic reprogramming in ECs following d-flow, we conducted ex vivo ^13^C metabolic tracing experiments using ECs isolated from partially ligated carotid arteries. PLCL was performed in both tamoxifen-treated non-transgenic littermate control (NLC) mice and *Lats1*<u>^+/−^</u>/*Lats2*^-/-^-EKO mice. Seven days after PLCL, ECs were isolated separately from the left carotid artery (LCA, exposed to disturbed flow) and the right carotid artery (RCA, exposed to laminar flow). Immediately after isolation, ECs were plated and allowed to adhere. At 24 hours post-plating, the culture medium was supplemented with either ^13^C-labeled glucose or 13C-labeled glutamine. At 48 hours post-plating, ECs were collected, metabolites were extracted, and carbon flux analysis was performed using mass spectrometry. This ex vivo approach enabled us to assess the incorporation of ^13^C-labeled substrates into key metabolic intermediates and compare the metabolic activity of ECs exposed to disturbed flow versus laminar flow in both genotypes. This analysis quantified the incorporation of ^13^C-labeled glutamine (A, B, F) or glucose (A, C, G) into various metabolic intermediates. The sizes of the circles in the diagrams represent the pool sizes of individual metabolites, while the color coding indicates the number of carbons incorporated from the ^13^C-labeled substrates (glutamine in (B, F) or glucose in (C, G)). The background arrows highlight the inferred major metabolic flows in ECs. The color gradient from gray to dark red reflects an increasing number of ^13^C-labeled carbons in the metabolites. **(D)** HUVECs transfected with siCont or siLATS1/2 were exposed to d-flow for 24 hours, and pyruvate dehydrogenase (PDH) activity levels were quantified. **(E)** HUVECs transfected with siCont or siLATS1/2, pre-treated with either the CD38 inhibitor 78c (1 µM) or vehicle, were exposed to d-flow, and protein expression was analyzed via Western blot. **(H, I)** HUVECs transfected with siCont or siLATS1/2 (H) or siRNA targeting SUOX (siSUOX) (I) were exposed to d-flow for 24 hours, and glutamate levels were measured. **(J)** HUVECs transfected with siCont or siSUOX were exposed to d-flow for 24 hours, and OGDH activity was measured as described in Methods. **(K)** HUVECs transfected with siCont or siSUOX for 48 hrs and treated with vehicle and MitoTEMPOL (10 µM) for one hour, then ECs were exposed to d-flow for 24 hours, and succinate was measured as described in Methods. **(L)** HUVECs transfected with siCont or siSUOX were exposed to d-flow for 24 hours, and complex V activity was measured as described in Methods. **(M)** HUVECs transfected with siCont or siSUOX for 48 hrs and treated with vehicle and oligomycine A (100 ng/ml)for one hour, then ECs were exposed to d-flow for 24 hours, and ATP was measured as described in Methods. **(N)** HUVECs transfected with siCont or siSUOX were exposed to d-flow for 24 hours, and SDH activity was measured as described in Methods. **(O)** HUVECs transfected with siCont or siLATS1/2, and after 48 hrs of transfection, ECs were pretreated with vehicle or BTB06584 (200 nM, specific inhibitor of ATP hydrolytic activity of mitochondrial FoF1-ATP synthase (complex V), without interfering with its ATP synthesis function) for one hour, and were exposed to d-flow for 24 hours, and p53, p53BP1, ICAM1, and Lamin B1 expression were detected by Western blotting (top). The percentages of SA-β-gal^+^ cells (bottom) were detected by flow cytometry. Quantification data for panels E and O are provided in the corresponding figure panels in Fig. S6. Data are presented as mean ± SD. Statistical tests, sample sizes, and detailed results for all panels are summarized in Table S1.

To compare the incorporation of [¹³C]glucose or [¹³C]glutamine into metabolic intermediates between NLC mice and *Lats1*^+/−^/*Lats2*^-/-^-EKO mice, we analyzed ECs isolated from the right carotid artery (RCA) exposed to l-flow. No significant differences in ^13^C incorporation into any metabolites were observed between the two genotypes under l-flow conditions (**Fig. S6A, B**). In contrast, under d-flow conditions in the LCA, we found that the reduced incorporation of ^13^C- labeled glucose and glutamine into TCA cycle intermediates observed in NLC mice was completely reversed in *Lats1*^+/−^/*Lats2*^-/-^-EKO mice (**Fig. 6F, G**). This reversal could not be attributed to changes in the expression or activity of OGDH or PDH, which were downregulated by both d-flow and LATS1/2 depletion (**Fig. 6D, E, J, S6C**). Instead, we observed a significant increase in glutamate levels associated with LATS1/2 and SUOX depletion under d-flow (**Fig. 6H, I**). Together with data showing that LATS1/2-mediated downregulation of SUOX reduces ATP production (**Fig. 5K**)—likely due to impaired electron transport chain (ETC) function— these findings suggest a compensatory upregulation of glutamate and TCA cycle intermediates upstream of the dysfunctional pathway of ETC (**Fig. 5L**).

Supporting this, previous reports have shown that the conversion of α-ketoglutarate (α-KG) to succinate via OGDH can be bypassed through a non-enzymatic reaction under conditions of elevated ROS^31^. Based on this, we hypothesize that the mtROS generated by d-flow and LATS1/2 deficiency—via SUOX downregulation (**Fig. 5L**)—may drive this non-enzymatic conversion. Indeed, treatment with the mtROS scavenger mitoTEMPOL significantly suppressed the d-flow- and siSUOX-induced succinate accumulation, suggesting that an mtROS-mediated, non-enzymatic pathway contributes to succinate production in the context of LATS1/2 deficiency, even though OGDH activity remains downregulated (**Fig. 6K**). Furthermore, as SUOX deficiency has been reported to cause complex V dysfunction through sulfite toxicity^30^, we assessed complex V activity. We found that both d-flow and SUOX depletion markedly inhibited complex V function (**Fig. 6L**).

While oligomycin A is a well-established inhibitor of ATP synthase (Complex V), its effects under d-flow and SUOX knockdown conditions appear to extend beyond simple inhibition. Rather than exacerbating ATP depletion, oligomycin A treatment unexpectedly increased ATP levels (**Fig. 6M**), suggesting that Complex V may be operating in reverse mode. In this reversed state, Complex V hydrolyzes ATP to maintain mitochondrial membrane potential (ΔΨₘ) when the proton gradient collapses due to impaired electron transport chain (ETC) function^32^. Blocking this reverse activity with oligomycin A likely prevents ATP consumption, thereby preserving intracellular ATP. Supporting this hypothesis, we observed an increase in succinate dehydrogenase (SDH) activity (**Fig. 6N**), indicating a compensatory upregulation of Complex II aimed at restoring electron flow and maintaining mitochondrial function. To further validate the role of reverse-mode Complex V activity in d-flow-induced SASP, we employed BTB06584, a compound reported to selectively inhibit the reverse activity of Complex V^33^. BTB06584 treatment effectively reversed hallmark SASP markers induced by d-flow and LATS1/2 depletion, including elevated SA-β-gal activity, reduced Lamin B1, and increased p53, p53BP1, and ICAM-1 expression (**Fig. 6O**). These findings highlight reverse-mode Complex V as a critical driver of SASP-related events under conditions of mechanical stress and Hippo pathway suppression. Moreover, our data suggest that this shift in mitochondrial function promotes enhanced [¹³C]glutamine incorporation into TCA cycle intermediates via upregulated SDH activity^34^, pointing to a broader metabolic reprogramming where glutamate utilization and upstream TCA metabolism are enhanced despite an overall reduction in ATP output (**Fig. 5L**).

### CD38 activation promotes d-flow-induced SAS In Vivo

While CD38 is traditionally recognized for its role in NAD⁺ depletion and the promotion of cellular senescence, we observed a positive correlation between CD38 and Ki67 expression in atherosclerotic plaques from *Lats1*^+/−^/*Lats2*^-/-^-EKO mice, suggesting that CD38 may contribute to d-flow-mediated induction of senescence-associated stemness (SAS) (**Fig. 3B**). Due to the challenges in assessing SAS in vitro^21^, we isolated ECs from carotid arteries following PLCL to evaluate SAS in vivo. ECs from *Lats1*^+/−^/*Lats2*^-/-^-EKO mice exhibited a higher proportion of SA-β-gal⁺/Ki67⁺ cells, confirming SAS induction. To assess the functional role of CD38, we treated mice with the CD38 inhibitor Ab68 or an IgG2 control prior to PLCL. Seven days post-ligation, ECs isolated from the left (LCA) and right (RCA) carotid arteries showed that CD38 inhibition markedly reduced SAS and restored the expression of LATS1/2, SUOX, and PDH in vivo (**Fig. 8A-C**). Taken together, these findings demonstrate that CD38 contributes to d-flow- and LATS1/2-deficiency-induced endothelial SAS by promoting metabolic reprogramming (**Fig. 5L**).

### The higher induction of senescence with metabolically active ECs in the human thrombogenic plaque compared to those in non-thrombogenic plaque

To investigate the roles of LATS1 and CD38 in human atherosclerotic plaques, we analyzed vascular specimens obtained from patients undergoing surgical repair of aortic aneurysms. Control (non-plaque) vascular samples were selected from donors or recipients in the lung and heart transplantation programs at Baylor Medical College (total *n* = 6; 3 males, 3 females; mean age 54.7 ± 11.7 years), while plaque-containing samples were selected from tissues obtained during aneurysm repair surgeries (total *n* = 8; 4 males, 4 females; mean age 68.6 ± 3.46 years). Using tissue microarray techniques, we prepared 15 tissue cores. Hematoxylin and eosin staining identified 7 cores without plaque lesions and 8 with definitive atherosclerotic lesions. We then performed IMC/ COMET™ analysis and integrated the data using the VISIOPHARM platform. Machine learning, trained on α-SMA staining patterns and cellular features, enabled precise compartmental mapping. In non-plaque samples, we identified surface, internal hyperplasia, media, and adventitia regions. In atherosclerotic samples, we defined surface, fibrous cap, plaque core, media, and adventitia areas (**Fig. 7D**). Additionally, a subset of plaques exhibited markedly increased both fibrin and TF staining, indicating a thrombogenic plaque (**Fig. S7B**).

**Figure 7.**
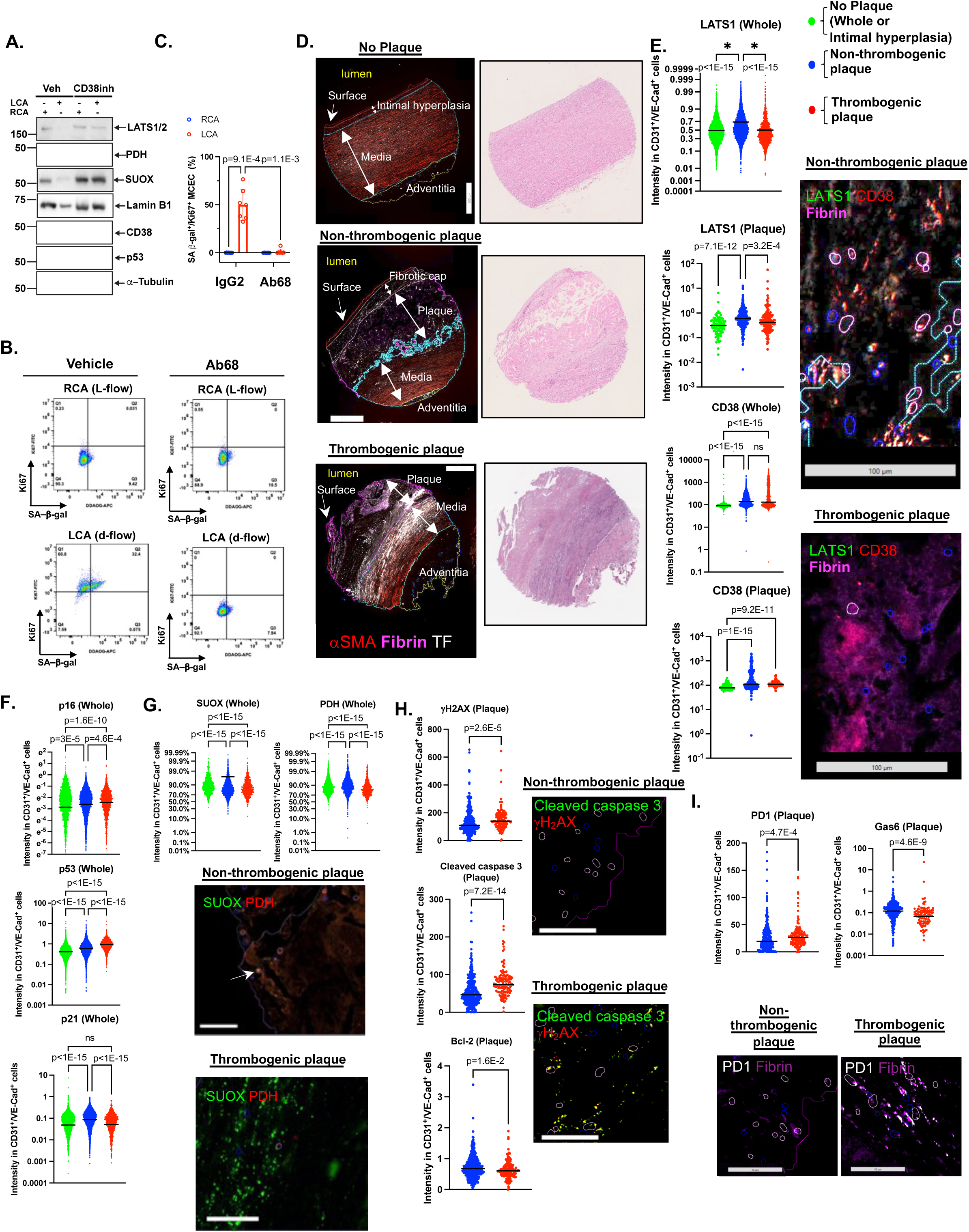
Inhibiting CD38 mitigates the downregulation of LATS1/2, SUOX, and PDH caused by d-flow, while also attenuating the upregulation of CD38 expression, SAS induction, and the formation of atherothrombosis. **(A-C)** 10-12-week-old WT C57BL/6 mice were injected with the CD38 inhibitor 78c (30 mg/kg/day), and PLCL was performed. Seven days post-ligation, LCA, exposed to d-flow, and RCA, exposed to l-flow, were perfused with an enzyme-containing solution to isolate ECs, as described in the Methods. Protein expression in the isolated ECs was analyzed via Western blot using the indicated antibodies (A). Co-expression of the fluorescent SA-β-gal marker and Ki67 was assessed in ECs isolated from the LCA and RCA following PLCL (B). The percentage of double-positive SA-β-gal and Ki67 cells was quantified and is displayed in the graph (C). **(D)** IMC and COMET™ analyses were performed on human atherosclerotic and no plaque lesions, with data integrated using the VISIOPHARM platform. Using machine learning trained on α-SMA staining and cellular morphology, we identified distinct anatomical regions: surface, internal hyperplasia, media, and adventitia in non-plaque lesions, and surface, fibrous cap, plaque core, media, and adventitia in plaque lesions. A subset of plaques showed high fibrin and tissue factor (TF) staining (**Fig. S7B**), indicative of thrombogenic plaques. **(E)** Expression levels of LATS1 and CD38 in double-positive CD31⁺/VE-Cadherin⁺ ECs selected from either the entire lesion area (Whole) or specifically from the plaque region (Plaque), which includes both the fibrous cap and plaque core as defined in (D). The intimal hyperplasia region from non-plaque lesions was used as a control to assess the phenotype of ECs within plaque areas. (**F, G**) Expression levels of senescence markers of p16, p53, and p21 (F), and SUOX and PDH expression (G) in double-positive CD31⁺/VE-Cadherin⁺ (ECs) selected from the entire lesion area (Whole) from no plaque (green), non-thrombogenic plaques (blue) and thrombogenic plaques (red). **(H, I)** Expression levels of the DNA damage marker γH2AX, apoptosis markers cleaved caspase-3 and Bcl-2 (H), as well as the “don’t eat me” signal PD-1 and the “eat me” signal Gas6 (I), in double-positive CD31⁺/VE-Cadherin⁺ ECs selected from the plaque regions of non-thrombogenic plaques (blue) and plaques with intraplaque hemorrhage or organized thrombosis (red). Statistical tests, sample sizes, and detailed results for all panels are summarized in Table S1.

We identified ECs by selecting double-positive cells expressing CD31 and VE-cadherin (VE- Cad), followed by cell segmentation analysis. To compare atherosclerotic plaques with non-plaque regions, we used the area of intimal hyperplasia in non-plaque lesions as a control, given its relatively higher EC content compared to other regions in no plaque samples. ECs from thrombogenic plaques showed markedly lower LATS1 expression in both the overall plaque area (whole) and the plaque core compared to ECs from non-thrombogenic plaques. CD38 expression was higher in ECs from thrombogenic plaques relative to no plaque regions; however, this increase was not significantly different when compared to ECs from non-thrombogenic plaques (**Fig. 7E**). Analysis of senescence markers revealed a significant upregulation of p16, p53, and p21 in ECs from non-thrombogenic plaques relative to no plaque lesions. Furthermore, p16 and p53 expression was even more elevated in ECs from thrombogenic plaques (**Fig. 7F**). On the other hand, SUOX and PDH expression were decreased in ECs from non-thrombogenic plaques compared to no plaque controls. (**Fig. 7G**).

Importantly, ECs from thrombogenic plaques exhibited significant increases in DNA damage marker γH2AX and the apoptotic marker cleaved caspase-3, along with a reduction in the anti-apoptotic protein Bcl-2 (**Fig. 7H**). Although efferocytosis function was not directly tested, we observed altered expression of related molecules in these ECs, including increased PD-1 and decreased Gas6 levels, which may impair efferocytosis by reducing pro-engulfment signaling and enhancing anti-engulfment cues, compared to ECs from non-thrombogenic plaques (**Fig. 7I**). Taken together, these data suggest the accumulation of senescent and apoptotic ECs in the plaque core, especially in the thrombogenic plaques. In these thrombogenic plaques, the expression of LATS1 was found to be lower than in plaques that are not thrombogenic, while the expression of CD38 was higher compared to areas without plaques.

We conducted spatial metabolomics analysis at a resolution of 20 µm and detected a total of 149 metabolites. By integrating these data with spatial proteomics and performing cell segmentation analysis, we were able to quantify metabolite levels in ECs with approximately single-cell resolution. We observed the presence of hydrogen sulfite (HSO₃⁻) and taurine, both metabolites linked to sulfite oxidase (SUOX) deficiency. HSO₃⁻ levels were significantly elevated in ECs within thrombogenic plaques compared to those in non-thrombogenic plaques (**Fig. 8A, B**). Interestingly, a pronounced increase in HSO₃⁻ was also detected specifically in the intimal hyperplasia regions of areas without plaque. This localized elevation may reflect increased hydrogen sulfide (H₂S) secretion from surface ECs in healthy, non-plaque regions^35^ (**Fig. 8A, B**). These findings are consistent with the observed downregulation of SUOX expression in ECs from thrombogenic plaques relative to ECs from non-thrombogenic plaques (**Fig. 7G**), suggesting a potential alteration in sulfur metabolism associated with thrombogenicity.

**Figure 8.**
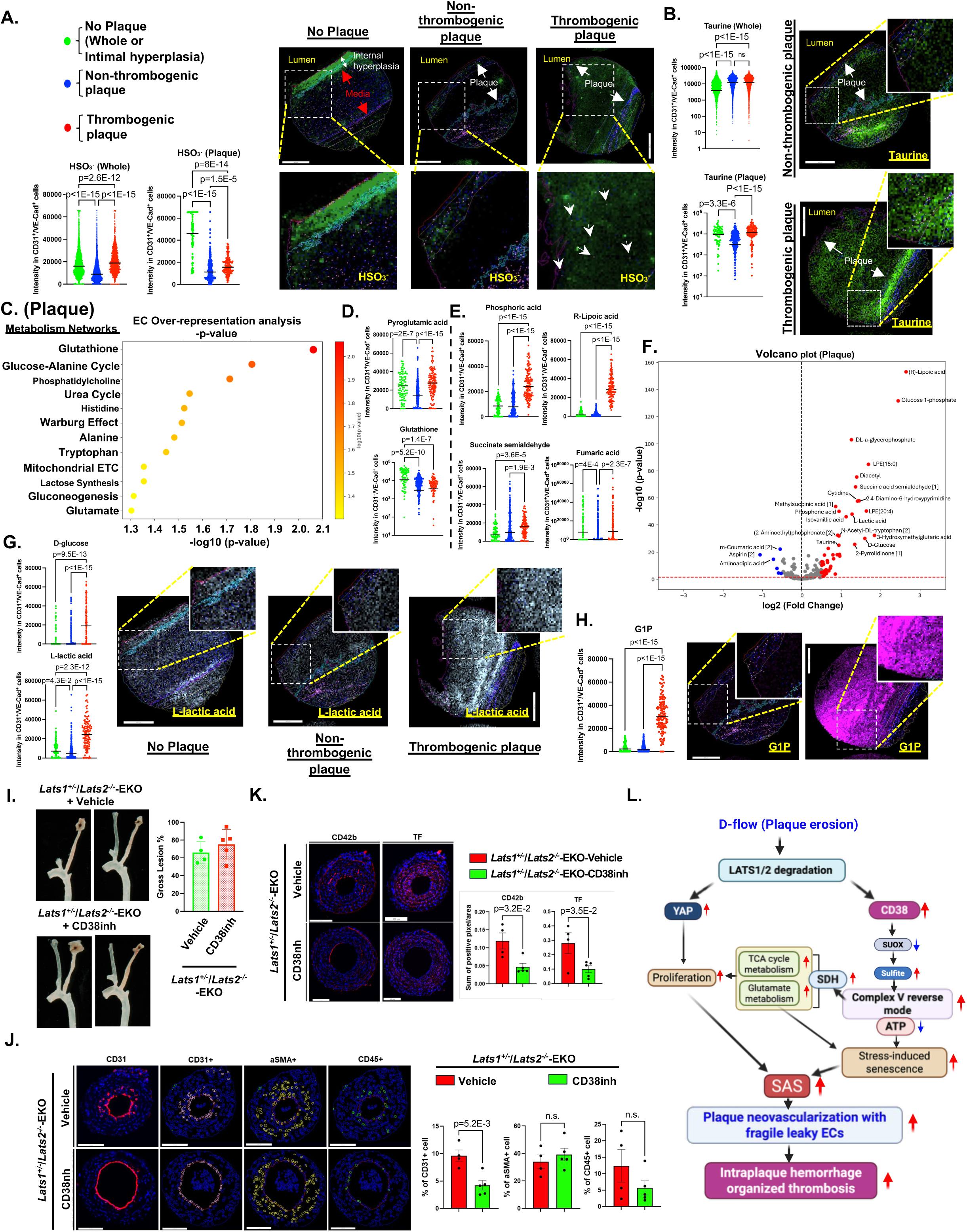
The upregulation of SUOX deficiency-related metabolites of hydrogen sulfite (HSO^3-^) and taurine, and glutamate, TCA cycle, and glycolysis -related metabolites in atherothrombotic plaque. (**A, B**) Expression levels of HSO3^-^ and taurine in double-positive CD31⁺/VE-Cadherin⁺ ECs selected from either the entire lesion area (Whole) or specifically from the plaque region (Plaque) in samples classified as no plaque (green), non-thrombogenic plaque (blue), and thrombogenic plaques (red). (**C**) By integrating IMC/COMET™ and spatial metabolomics data, EC over-representative analysis was performed, and single-cell metabolite profiles were obtained in double-positive CD31⁺/VE-Cadherin⁺ ECs selected specifically from the plaque region (Plaque) in samples classified as no plaque (green), non-thrombogenic plaque (blue), and thrombogenic plaque (red). (**D, E**) Expression levels of (D) pyroglutamic acid and glutathione, both involved in glutamate utilization and recycling through the glutathione cycle, and (E) phosphoric acid (which supports energy-dependent processes in glutamate and TCA cycle metabolism), succinate semialdehyde (a GABA shunt intermediate linked to glutamate metabolism), R-lipoic acid (a cofactor essential for the activity of PDH and OGDH, thereby supporting both glutamate metabolism and TCA cycle entry), and fumaric acid (a core TCA cycle intermediate), in double-positive CD31⁺/VE-Cadherin⁺ ECs selected specifically from the plaque region (Plaque) in samples categorized as no plaque (green), non-thrombogenic plaque (blue), and thrombogenic plaque (red). **(F)** Volcano plot illustrating the 149 metabolites detected in ECs from spatial metabolomics analysis of human plaques from plaques with intraplaque hemorrhage/organized thrombosis and non-thrombosed plaque. (**G, H**) Expression levels of (G) D-glucose and L-lactic acid, key metabolites in glycolysis, and (H) glucose-1-phosphate (G1P), an intermediate primarily involved in glycogen metabolism and indirectly linked to glycolysis, in double-positive CD31⁺/VE-Cadherin⁺ endothelial cells (ECs) selected specifically from the plaque region (Plaque) in samples classified as no plaque (green), non-thrombogenic plaque (blue), and thrombogenic plaque (red). **(I-K)** NLC and *Lats1***<u>^+/−^</u>**/*Lats2*^-/-^-EKO mice were administered the CD38 inhibitor 78c (CD38inh) (30 mg/kg/day) starting one day prior to PLCL. Atherosclerosis formation was evaluated two weeks post-ligation, as detailed in Fig. 3A. Gross plaque size was measured in NLC and *Lats1***<u>^+/−^</u>**/*Lats2*^-/-^-EKO mice after four weeks of PLCL (E). **(J)** COMET™ imaging followed by automated segmentation and quantification of CD31⁺ ECs,, αSMA⁺ VMSCs, and CD45⁺ leukocytes in LCA plaques. (right) Quantification of individual cell populations (CD31⁺, αSMA⁺, and CD45⁺) in vehicle and CD38inh-treated *Lats1*^+/−^/*Lats2*^-/-^-EKO plaques. EC count was significantly inhibited in CD38inh-treated *Lats1*^+/−^/*Lats2*^-/-^-EKO lesions comopared to vehicle-treated ones (n = 4–5/group; mean ± SD). (**K**) Representative immunofluorescence staining of CD42b and TF in plaques from vehicle and CD38inh-treated *Lats1*^+/−^/*Lats2*^-/-^-EKO mice. Decreased deposition of CD42b and TF is observed in CD38inh-treated *Lats1*^+/−^/*Lats2*^-/-^-EKO mice lesions compared to those in vehicle-treated ones. Data are presented as mean ± SD, n=4-5. The applied statistical tests, sample number, and results in all figures are summarized in Table S1. **(L) Proposed model of D-flow–induced senescence-associated stemness (SAS) driven by LATS1/2 downregulation:** Under d-flow, downregulation of LATS1/2 activates YAP, leading to upregulation of CD38. CD38 activation suppresses SUOX expression, resulting in sulfite accumulation. This disrupts mitochondrial redox homeostasis and promotes the reverse mode of mitochondrial complex V, altering bioenergetics and triggering upstream metabolic rewiring. One key consequence is the activation of succinate dehydrogenase (SDH), which enhances TCA cycle activity and glutamate metabolism. This metabolic reprogramming supports the biosynthesis of nucleotides and amino acids—critical precursors for DNA, RNA, and protein synthesis—thereby fueling EC proliferation and pathological neoangiogenesis. However, the shift to reverse complex V function leads to reduced ATP production, imposing metabolic stress on ECs. This stress, combined with enhanced TCA and glutamate pathway flux, promotes a SAS phenotype—characterized by excessive EC proliferation alongside features of cellular senescence. The resulting imbalance drives abnormal, fragile neovessel formation and contributes to atherothrombosis, distinct from the more stable lesions typically seen in standard mouse models of atherosclerosis.

Additionally, we performed EC over-representation analysis focusing on metabolites within the plaque area (**Fig. 8C**). This revealed significant metabolic alterations between non-thrombogenic and thrombogenic plaques. Specifically, differential enrichment was identified in glutathione metabolism, the glucose-alanine cycle, urea cycle, and Warburg effect-related pathways, implicating enhanced activity in glycolysis, the TCA cycle, and glutamate metabolism. Glutathione levels were lowered in both non-thrombogenic and thrombogenic atherosclerotic plaques compared to the internal hyperplasia region of no plaque lesions. In contrast, pyroglutamic acid was significantly elevated in thrombogenic plaques relative to non-thrombogenic ones, suggesting impaired glutathione recycling and increased oxidative stress in more advanced, thrombogenic lesions (**Fig. 8D**).

We also observed elevated levels of phosphoric acid, succinic semialdehyde, R-lipoic acid, and fumaric acid in ECs from thrombogenic plaques compared to non-thrombogenic ones (**Fig. 8E, F)**. These metabolites point to enhanced metabolic activity and mitochondrial stress. In particular, increased succinic semialdehyde suggests activation of the GABA shunt, a stress-responsive pathway linked to glutamate metabolism. Elevated R-lipoic acid, a key mitochondrial cofactor for enzymes like PDH and OGDH, may indicate a compensatory effort to sustain mitochondrial function and TCA cycle entry under energy stress. Higher fumaric acid levels, a central TCA cycle intermediate, further support increased flux or altered regulation of this pathway. Together, these changes suggest a metabolic shift in ECs within thrombogenic plaques toward alternative energy pathways and redox adaptation, aligning with the metabolic phenotype observed in Lats1⁺/⁻/Lats2⁻/⁻-EKO mice (**Fig. 5, 6**). Consistently, multiple metabolites involved in glycolysis, the TCA cycle, and glutamate metabolism were significantly upregulated in ECs from thrombogenic plaques (**Fig. 8F–H**). Despite signs of senescence and apoptosis (**Fig. 7H**), these ECs remain metabolically active, reflecting a complex and adaptive metabolic state.

Notably, we identified a striking accumulation of hydrogen sulfite (HSO₃⁻) and taurine— metabolites typically associated with sulfite oxidase (SUOX) deficiency. HSO₃⁻ levels were significantly higher in ECs from thrombogenic plaques compared to non-thrombogenic plaque (**Fig. 8A**), in parallel with a marked downregulation of SUOX expression (**Fig. 7G**). This previously unrecognized alteration in sulfur metabolism highlights a potentially novel link between SUOX deficiency and endothelial dysfunction in atherothrombosis, offering new insight into disease progression and therapeutic targets.

## DISCUSSION

Our research used *Lats1*^-/-^/*Lats2*^-/-^-EKO mice to show significant systemic edema caused by disrupting the EC barrier. Additionally, creating EC-specific *Lats1*^+/−^/*Lats2*^-/-^-EKO mice led to atherothrombosis, characterized by increased neoangiogenesis, leukocyte infiltration, and decreased α-SMA positive cells under hypercholesterolemia and d-flow. These results suggest the development of advanced plaques similar to those seen in humans. To understand how endothelial LATS1/2 contributes to neoangiogenesis, thrombosis, inflammation, and senescence—all factors in atherothrombosis—we used the IMC/COMET™ method. This technique allows for qualitative in vivo analysis at the single-cell level of these cellular phenotypes. We observed an upregulation of markers related to proliferation (Ki67 and YAP), thrombosis (TF, CD42b, and fibrin), and inflammation (CCR7 and IL-6). We also noted a significant increase in senescence markers p53 and CD38 in the ECs of *Lats1*^+/−^/*Lats2*^-/-^-EKO mice. Our study identifies a non-canonical function of CD38 in ECs subjected to d-flow. Under physiological conditions, CD38 is a well-established marker of cellular senescence, typically linked to irreversible cell cycle arrest and loss of proliferative potential (Group 1 and 3 in Fig. 3B). However, in LATS1/2-deficient ECs, we observed a distinct population characterized by simultaneously high CD38 and Ki67 expression, with a linear correlation between the two markers (SAS, Group 2 in Fig.3B). This indicates that these cells retain senescent features while escaping growth arrest, giving rise to a paradoxical state where senescence and proliferation coexist. CD38 thus marks a unique endothelial phenotype capable of contributing to neoangiogenesis in atherosclerotic lesions. These findings challenge the classical view of senescence as a strictly anti-proliferative state and underscore a context-dependent shift in CD38’s functional role. SAS ECs contribute not only to increased plaque neoangiogenesis, but also to the accumulation of senescent cells and the development of fragile, leaky microvessels, hallmarks of advanced atherosclerotic lesions (**Fig. 8L**).

Plaque neovascularization, or neoangiogenesis, involves the formation of new blood vessels within atherosclerotic plaques. These new vessels are often fragile and prone to rupture, leading to intraplaque hemorrhage^36^. This hemorrhage can contribute to plaque instability and increase the risk of plaque rupture. When a plaque ruptures, it exposes thrombogenic material to the bloodstream, triggering the formation of a blood clot, or thrombus, which can obstruct blood flow and lead to atherothrombosis^37,38^. Thus, neovascularization within plaques plays a critical role in the progression of atherosclerosis and the development of atherothrombosis, linking these processes to increased cardiovascular risk^36^. Our study demonstrated that the reduction of LATS1/2, which mediates CD38 activation, is a key factor in triggering plaque neoangiogenesis. This process is characterized by the formation of leaky and fragile vessels, driven by endothelial cell SAS, ultimately resulting in atherothrombosis (**Fig. 8L**).

SUOX is a mitochondrial enzyme responsible for converting toxic sulfites into harmless sulfates, which are excreted from the body. A deficiency in SUOX, typically caused by genetic mutations in the SUOX gene, leads to the accumulation of sulfites in tissues, resulting in severe metabolic and neurological complications. This autosomal recessive condition manifests with symptoms such as seizures, intellectual disability, developmental delays, elevated sulfite and taurine levels in urine, feeding difficulties, hypotonia, and progressive neurodegeneration^28,29^. Our research also revealed that SUOX deficiency, induced by d-flow and LATS1/2 depletion, decreased complex V-ATP synthase activity, leading to decreased ATP levels.

Notably, while d-flow suppressed the enzymatic activities of OGDH and PDH in ECs from NLC mice, this inhibition was bypassed in *Lats1*^+/−^/*Lats2*^-/-^-EKO mice via ROS-mediated non-enzymatic pathways, particularly under conditions of SUOX deficiency. Additionally, we found that LATS1/2 depletion led to a reduction in SUOX expression, which triggered the reverse mode of mitochondrial complex V. This shift activated SDH, resulting in increased ATP consumption. This bypass mechanism, involving OGDH and SDH activation, enhanced the incorporation of glutamine and glucose into TCA cycle intermediates and led to a compensatory response to elevated ATP demand at the ETC. Consequently, there was a marked upregulation of upstream metabolic pathways—including glycolysis, glutamate metabolism, and the TCA cycle. This shift enhances glutamate metabolism, known to induce cellular senescence through ammonium (NH₄⁺) accumulation^39^, and stimulates TCA cycle activity despite a reduction in ATP production. These metabolic shifts also provided essential precursors for DNA, RNA^40^, and protein synthesis^41,42^, thereby supporting increased EC proliferation and neoangiogenesis within atherosclerotic plaques. However, despite these anabolic adaptations, ATP synthesis remained impaired, triggering a stress-induced senescence response. Thus, while LATS1/2 depletion facilitates neovascularization even under ATP-deficient conditions, it promotes the formation of fragile and leaky microvessels, ultimately contributing to intraplaque hemorrhage and organized thrombosis (**Fig. 8L**).

To support these findings, we observed significantly distinct senescence and metabolic profiles in ECs from non-thrombogenic plaques compared to thrombogenic ones. In thrombogenic plaques, ECs showed increased accumulation of senescent and apoptotic cells, accompanied by lower expression of LATS1, SUOX, and PDH, and elevated expression of CD38. This pattern underscores the involvement of the LATS1/2–CD38 axis in promoting EC senescence and apoptosis. Moreover, spatial metabolomics analysis revealed that ECs within thrombogenic plaques were metabolically hyperactive, with marked upregulation of metabolites associated with glycolysis, the TCA cycle, and glutamate metabolism. These observations suggest the presence of metabolically active senescent ECs in human thrombogenic plaques, mirroring the phenotype observed in *Lats1*^+/−^/*Lats2*^-/-^-EKO mice. This metabolic reprogramming was associated with a distinct EC phenotype characterized by heightened SAS, which drove abnormal EC proliferation, premature senescence, and eventual cell death in areas of atherothrombosis. Collectively, these cellular and metabolic changes contributed to the progression of atherothrombosis. Notably, pharmacological inhibition of CD38 effectively mitigated these pathological outcomes, highlighting its potential as a therapeutic target.

A key distinction between human atherothrombotic lesions and those observed in *Lats1*^+/−^/*Lats2*^-/-^ -EKO mice lies in the extent of angiogenesis. While we observed EC neovascularization within human plaques, there was no significant overall increase in angiogenesis. This likely reflects the chronic nature of atherothrombotic plaque development in humans, where the local microenvironment is harsh and unfavorable for sustained EC survival. As a result, extensive EC death may limit the expansion of EC populations, masking any apparent increase in angiogenesis despite neovascular activity. Another important consideration is the origin of our human atherosclerotic samples. All specimens were obtained from patients undergoing elective surgery for aortic aneurysm repair. Consequently, these samples likely represent a chronic or stabilized phase of atherothrombosis, rather than an acute event such as plaque rupture. In this context, the observed intraplaque hemorrhage and organized thrombosis may not result solely from leaky neovessels but also from repetitive thrombus deposition and organization. These surface thrombi could arise from minor erosions, undergo partial resolution, and become incorporated into the plaque structure, ultimately forming intraplaque hemorrhages or organized thrombotic regions. Given these findings, we propose that d-flow-mediated decrease of LATS1/2 may play a critical role in promoting surface erosion–driven repetitive thrombus deposition. This mechanism may preferentially affect ECs localized to the luminal surface of the plaque. Future studies should address these two possibilities—(1) leaky neovessels and (2) recurrent surface thrombosis—as potential contributors to the development of intraplaque hemorrhage and organized thrombotic lesions in human atherosclerosis.

In conclusion, LATS1/2 deficiency induced by d-flow promotes the development of a SAS phenotype in ECs, characterized by a paradoxical combination of cellular senescence and regenerative potential. This phenotype contributes to the formation of fragile, leaky neovessels— hallmarks of advanced human atherothrombotic lesions. Both YAP activation and CD38 play key roles in establishing SAS. Specifically, CD38-mediated depletion of SUOX induces the reverse mode of mitochondrial complex V, activating SDH and resulting in ATP depletion. This energy stress triggers compensatory upregulation of the TCA cycle and glutamate metabolism, providing biosynthetic support for the SAS phenotype. The induction of SAS through LATS1/2 degradation and CD38 activation supports EC proliferation and neoangiogenesis even under metabolically adverse, ATP-deficient conditions. However, the resulting neovessels are structurally compromised, contributing to intraplaque hemorrhage and organized thrombosis.

This mechanism is prominent in human atherothrombotic lesions but not recapitulated in standard hypercholesterolemic mouse models, highlighting the translational importance of LATS1/2–CD38–SAS signaling in human vascular pathology. These findings position CD38 as a promising therapeutic target for mitigating atherothrombotic progression.

## Acknowledgements

We thank Scientific Publications, Research Medical Library at The University of Texas MD Anderson Cancer Center for editing and Carolyn J. Giancursio for her technical assistance.

## Contributions

S.K. and J.L. performed the experiments, interpreted the data, and wrote the manuscript. K.A.K. performed the experiments and analyzed the data. W.C. supported data analyses, and wrote the manuscript. Y.J.G., V.K.S. and O.H. performed the experiments and maintained mouse colonies. G.F.M performed the surgery and the following analysis. S.L. supported data analysis. K.L.S, L.A.R.,A.D., J.P.C. K.F., N.L.P., E.K., and J.H. contributed to the interpretation of the data. H.G.V. and Y.H.S. provided the human atherosclerosis samples and contributed to the interpretation of the data. K.C.T.S., J.H.K., K.C.O.M. performed histological analysis. R.P. supported animal surgery and contributed the interpretation of the data. P.L.L., L.T., I.M. performed metabolite assay and contributed the interpretation of the data. L.YC and P.S. supported metabolite analysis and contributed the interpretation of the data. C.M. supported the thrombosis analysis. E N.C provided CD38 inhibitor and supported the data analysis. J.F.M. provided LATS1 and LATS2 flox/flox mice, and contributed the interpretation of the data. H.X., supported the histological analysis and diagnosis. E.H.S performed the mass spectrometry imaging and aided in its data interpretation. J.K.B. planned and analyzed imaging mass cytometry and COMET data. S.K., NT.L., G.W., and J.A. planned and generated the study design, obtained funding, interpreted data, and wrote the manuscript.

## Supplemental Materials

Supplementary Methods

Contact for Reagent and Resource Sharing

Experimental model and Subject Details

Methods Details

Online Supplementary Figures S1-S7

Online Supplementary Figure Legends S1-7

Online Supplementary Table 1-3

Raw Data Sets File1-6

## Non-standard Abbreviations and Acronyms

53BP1: p53-binding protein 1
AAV8-PCSK9: adeno-associated-virus-8 overexpressing pro-protein convertase subtilisin/Kexin type 9 gain-of-function D377Y mutant
d-flow: disturbed flow
ECs: endothelial cells
EKO: endothelial-specific knockout
HFD: high-fat diet
IMC: imaging mass cytometry
Ki67: marker of proliferation Ki-67
LATS1/2: Large Tumor Suppressor Kinase 1and 2
LCA: Left common carotid artery
mtROS: mitochondrial reactive oxygen species
NAD^+^: nicotinamide adenine dinucleotide
NCD: normal chow diet
NLC: Non-knockout littermate controls
p53: tumor protein p53
p16: cyclin-dependent kinase inhibitor 2A
p21: cyclin-dependent kinase inhibitor 1
PDH: pyruvate dehydrogenase
PLCL: Partial left carotid ligation
RCA: Right common carotid artery
ROS: reactive oxygen species
ROI: region of interest
S-Βgal: Senescence-Associated β-Galactosidase
SAS: senescence-associated stemness
SASP: senescence-associated secretory phenotype
SDH: succinate dehydrogenase
SUOX: sulfite oxidase
TRX1: thioredoxin 1
YAP: Yes-associated protein

